# The Ptch methylase installs an m^3^U modification on 28S rRNA for efficient protein synthesis in flies and men

**DOI:** 10.1101/2024.06.27.600882

**Authors:** Jie Chen, Yaofu Bai, Yuantai Huang, Min Cui, Yiqing Wang, Zhenqi Gu, Xiaolong Wu, Yubin Li, Yikang S. Rong

## Abstract

The ribosomal RNA (rRNA) is one of the most heavily modified RNA species in nature. Although we have advanced knowledge of the sites, functions and the enzymology of many of the rRNA modifications from all kingdoms, we lack basic understanding on many of those that are not universally present. A single N^3^ modified Uridine base (m^3^U) was identified on the 28S rRNA from human and frog over thirty years ago, which is absent in bacteria or yeast. Here we show that the equivalent m^3^U is present in Drosophila, and that the Ptch enzyme and its human homolog are both necessary and sufficient for carrying out the modification. The Ptch-modified U is at a functional center of the large ribosome, and consistently *ptch*-mutant cells suffer loss of ribosomal functions. Ptch, proposed to be the most druggable RNA methyltransferases in human, represents a unique target where ribosomal functions could be specifically compromised in cancer cells.

## Introduction

Chemical modifications play important roles for the functions of RNA molecules. The RNA component of the Ribosome, ribosomal RNA (rRNA), is the most abundant RNA species in cells and has one of the most abundant modifications in both class and number (for recent reviews see Sloan et al. 2017; Sergiev et al. 2018; Londei and Ferreira-Cerca 2021; Streit and Schleiff 2021). The most abundant modifications on rRNA are 2’-O-methylation of the sugar moiety and pseudo-uridylation of the Uridine base. The biological functions of these modifications involve regulating RNA stability and base interactions between subunits or between different functional centers of the ribosome so to control translational efficiency and accuracy (reviewed in Helm 2006; Baxter-Roshek et al. 2007; Liu et al. 2008; Baudin-Baillieu et al. 2009; Mahto and Chow 2013; Erales et al. 2017). Many of these modifications are evolutionarily conserved. Most of the 2’-O-methylation and pseudo-uridylation are implemented by methylases, such as Fibrillarin, directed by small nucleolar RNAs (snoRNAs) that base-pair with rRNA at the targeted site (reviewed in Maxwell and Fournier 1995; Kiss-László et al. 1996). However, “stand-alone” methylases also exist for these modifications, such as the Pet56 enzyme from bacteria (Sirum-Connolly and Mason 1993) and Spb1 from yeast (Bonnerot et al. 2003; Lapeyre and Purushothaman 2004).

Methylation of bases on rRNA is also abundant, such as m^6^A, m^5^C and m^3^U. These modifications are carried out by “stand-alone” methylases that do not rely on snoRNAs (e.g., Sharma et al. 2014; Bourgeois et al. 2015; van Tran et al. 2019). Base methylation also carries essential functions for the ribosome. For example, the KsgA enzyme from bacteria installs specific m^6^A modification for the regulation of translational accuracy (van Buul et al. 1984; Pletnev et al. 2020). Similar to both 2’-O-methylation and pseudo-uridylation of rRNA, base methylation display species specific class and location, suggesting specialized functions.

Methylation at the N^3^ position of Uridine (m^3^U) is one of the least abundant base methylations in nature. The m^3^U base has a reduced base-pairing strength with the A base due to the loss of one of the two hydrogen bonds. In *E. coli*, the RsmE enzyme is responsible for the methylation of U1498 on 16S rRNA, the loss of which results in growth defects with a not-well understood mechanism (Basturea et al. 2006; Basturea and Deutscher 2007; Zhang et al. 2012). Two modified U bases have been identified on 25S rRNA from the budding yeast, and the enzymes Btm5 and Btm6 were identified as the responsible methylases (Sharma et al. 2014). Unfortunately, how these modifications contribute to ribosomal functions is also poorly understood.

In 1990, Maden reported a single N^3^ modified U base on human and frog rRNAs, which was further confirmed in later studies (Maden 1990; Taoka et al. 2018). The modified base of U4500 of the huamn 28S rRNA, with naming according to the 3D Ribosomal Modification Maps Database (Piekna-Przybylska et al. 2008), is one of the two neighboring and universally conserved U bases at the Peptidyl Transferase Center (PTC), an essential functional center of the large ribosome (reviewed in Polacek and Mankin 2005). Interestingly, that U base at PTC is not modified on bacteria or yeast rRNA, suggesting specialized function of this m^3^U in higher organisms.

Here we identify the enzyme responsible for this modification. By mutational analyses, we showed that the conserved Ptch protein is required for the presence of m^3^U at eukaryotic PTCs. In addition, purified Ptch proteins from flies and men are capable of methylating rRNA substrates *in vitro*. Furthermore, Drosophila cells without an m^3^U modified PTC suffer a growth defect accompanied by reduced protein synthesis, while loss of the enzyme is lethal to human cells. The identification of Ptch, a member of the SPOUT class of methyltransferases, as the m^3^U methylase of eukaryotic organisms helps fill an outstanding gap in our understanding of rRNA modifications.

## Results

### Ptch is a nucleolar protein essential for growth and viability

We conducted a small scale CRISPR/Cas9 mediated mutagenesis of Drosophila genes predicted to encode domains with potential RNA modifying activities (e.g. Peng et al. 2023). For the uncharacterized *CG12128* gene, we recovered frameshift mutations at the C-terminal region that cause recessive phenotypes, which include partial lethality of homozygotes (∼75%) and complete female sterility of the survivors while male homozygous mutants are semi-fertile (Figure S1). The genomic structure of *CG12128* and alleles used in this study are provided in Figure 1A. DAPI staining of the mutant ovaries revealed that they are filled with egg chambers arrested at early stages of oogenesis (Figure 1B). Based on the morphology of the mutant ovaries, we rename *CG12128* as *<u>p</u>u tao <u>ch</u>uan* (*ptch*), which is the Chinese translation of “strings of grapes”. In addition to semi-lethality and female sterility, *ptch* mutant adults display a classic “Minute” phenotype indicative of partial loss of ribosomal functions (Saebøe-Larssen et al. 1998; Marygold et al. 2007), which includes slow growth, small body sizes, thin and shortened bristles, and sterilities in adults (Figure 1C, Figure S1B).

**Figure 1.**
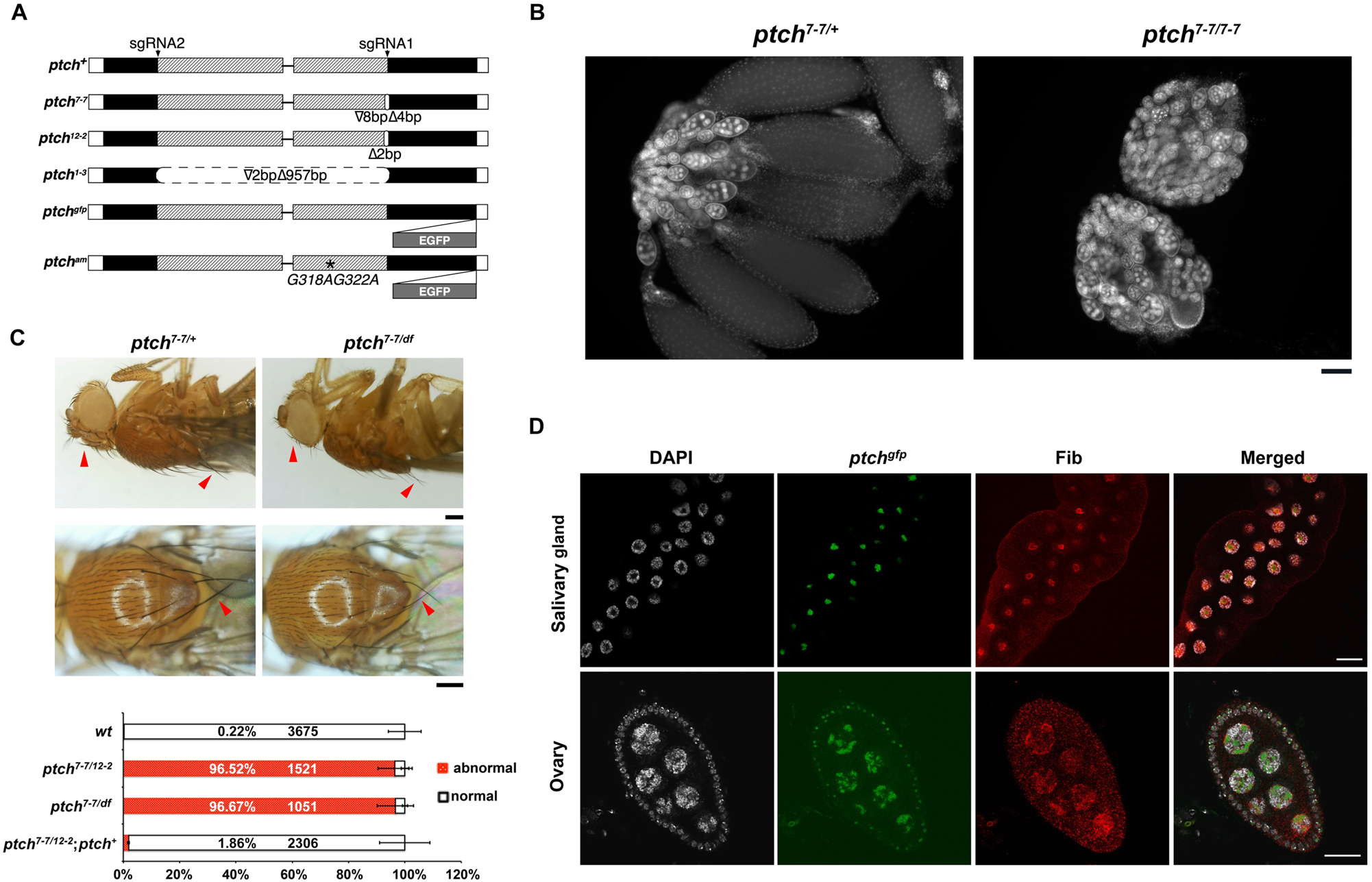
Ptch is a nucleolar protein essential for animal development. **A. Alleles of *ptch*.** Rectangular boxes represent exonic regions, and lines indicate introns. Unfilled (white) regions represent UTRs, and filled regions indicate coding regions. The hatched region encodes the methyltransferase domain. The two sgRNA-targeted regions are indicated by arrowheads. The approximate positions of the point mutations are indicated by vertical white bars. In the two transgenic rescuing constructs at the bottom, the EGFP tag and the *G318AG322A* mutation (am) are indicated. Drawings are not to scale. **B. “Minute” bristles on *ptch* mutants.** In the dorsal and the side views of adult flies, the largest bristles on the back and head regions are indicated with red arrowheads, showing the mutants having thinner and shorter bristles. Genotypes are listed on top of the pictures. Quantification of the bristle phenotypes are given below the pictures, with genotypes given to the left. *df*: a chromosomal deficiency of the *ptch* region. **C. Defective oogenesis of *ptch* mutants.** DAPI staining of dissected adult ovaries, with genotypes on top, shows normal development in wild type females in which less mature egg chambers are heavily stained for DAPI while mature eggs are less stained. In the mutants, only immature egg chambers are present. Scale bar represents 100μm. **D. Nucleolar localization of Ptch.** Immunostaining of salivary glands from third instar larvae and ovaries from adults, both with the genotype of *ptch^1-3/1-3^; [ptch^gfp/gfp^]*, were performed with an anti-Fib antibody. A single gland or a single egg chamber were shown with DNA signal in white, Ptch^GFP^ fluorescence signal in green and anti-Fib signal in red. Merged images of all three signals are also shown. Scale bars represent 40μm. For more Ptch localization images see Figure S1 in Supplemental materials.

We also generated a deletion mutation of *ptch* using two sgRNAs in CRISPR/Cas9 mutagenesis, deleting the entire methyltransferase domain (Figure 1A). This mutation, likely null, causes complete lethality at the third instar stage suggesting that the aforementioned frameshift mutations are likely hypomorphic in nature. All the phenotypes described above: lethality, sterility, “Minute”, were rescued with a transgene that carries a genomic fragment of the wildtype *ptch* locus. So was a similar transgene carrying *ptch* but with a C-terminally tagged *egfp* gene able to rescue the mutant phenotypes (Figure 1A). In addition, we constructed a rescuing transgene in which two Gly residues predicted to be essential for the methyltransferase activity are mutated to Ala (Ptch^G318AG322A^, Figure S1, Figure 1A). This mutated Ptch protein (Ptch^am^, active-site mutant), although able to correctly localize (see below and Figure S1), is unable to substitute for the function of the wildtype protein (Figure S1). These results taken together confirm that the mutant phenotypes are due to the loss of Ptch, and primarily the loss of its methylase function.

To determine the cellular location of Ptch, we first used GFP fluorescence in tissues of *ptch^gfp^* animals in which the only class of Ptch proteins are those tagged with EGFP (full genotype: *ptch^1-3/1-3^; ptch^gfp/gfp^*). We discovered that Ptch-GFP are concentrated in the nucleolus. This nucleolar localization was confirmed when Ptch-GFP signals were merged with anti-Fibrillarin (Fib) signals in immunostaining experiments, using Fib as a nucleolar marker (Figure 1D). Ptch also co-localizes with rRNA in the nucleolus in immuno-FISH experiments (Figure S1). Therefore, Ptch is a nucleolar protein, likely carrying out molecular functions related to rRNA, which is consistent with the Minute phenotypes described earlier. Interestingly, the Human Ptch homolog, C9orf114/SPOUT1, has been identified as essential for cell viability (Wang et al. 2015; Treiber et al. 2017), and overexpressed C9orf114 protein localizes to the nucleolus (Wang et al. 2015).

### The m^3^U modification of 28S rRNA depends on Ptch

The Ptch protein has an annotated methyltransferase domain belonging to the SPOUT family. Members of this family are RNA methyltransferases acting on nucleotide bases on either the rRNA or tRNA (Anantharaman et al. 2002; Tkaczuk et al. 2007; Hori 2017; Strassler et al. 2022). The nucleolar localization of Ptch led us to hypothesis that Ptch performs base methylation of rRNA.

A conventional way to detect some of the RNA modifications, particularly 2’-O-methylation, is based on reverse-transcriptase’s reduced ability to traverse a modified base on the RNA template in the presence of restrictive amounts of dNTPs. We employed the method of “Reverse Transcription at Low deoxy-ribonucleoside triphosphate concentrations followed by Polymerase chain reaction” (RTL-P, Dong et al. 2012) to sample a few regions of the rRNA, and identified a small region towards the 3’ end of 28S rRNA that yield different RT-PCR patterns between the *wt* and *ptch* mutants (Figure S2A, B). The region that we identified by RTL-P is highly conserved in eukaryotes (Figure S2C). RNA modifications on human 28S rRNA have been extensively categorized previously (Taoka et al. 2018). We noted that a single N^3^ modification of U4500 of human and U3629 of frog rRNAs was discovered over 30 years ago (Maiden 1990), and the responsible methyltransferase remained unknown.

To investigate whether the corresponding U base (U3485) in fly 28S rRNA is modified (Figure 2A), we employed a primer extension method based on that an m^3^U base partially losses hydrogen-bonding capacity with the A base (Figure 2B), leading to termination of reverse transcription. A similar assay has been used in m^3^U studies in bacteria and yeast (Basturea and Deutscher 2007; Sharma et al. 2014). As shown in Figure 2C, reversed transcription using total RNA from *wt* or *ptch*-heterozygous adults as an RT template produced fragments terminating at the base immediately 3’ to the potentially modified U base, confirming the existence of a modification. More importantly, the termination was eliminated when *ptch* homozygous mutant RNA was used in RT, indicating that the modification is abolished by the mutation. Similar primer extension confirms the presence of the modified U on rRNAs from human and mouse but its absence from either budding or fission yeast rRNAs (Figure 2D).

**Figure 2.**
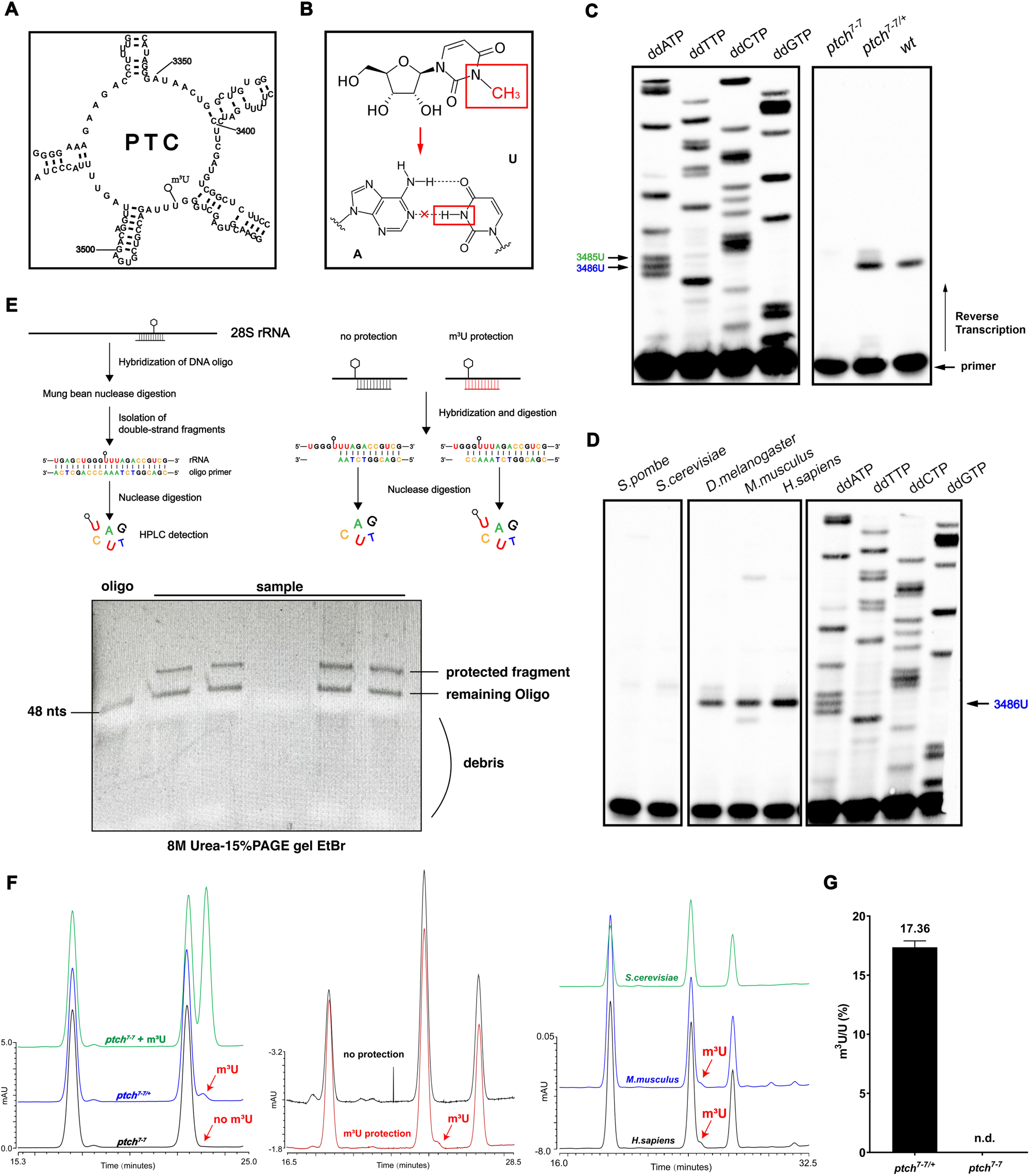
m^3^U modification of 28S rRNA is controlled by Ptch. **A. Schematic representation of PTC from Drosophila.** Secondary structure was reproduced from https://rnacentral.org/rna/URS0000A53741/7227, with base numbering and the m^3^U position labelled. **B. The chemical structure of m^3^U and its pairing with the A base.** At the top is the m^3^U with the methyl group outlined in red. At the bottom, the hydrogen bonds are shown for a normal A:U pair. The would-be methylated N in m^3^U is outlined in red. **C. Primer extension detection of m^3^U in Drosophila RNA.** At the right, total RNA from adults of different genotypes, listed on top of the gel picture, were subject to primer extension by reverse transcription. At the left is a sequencing gel, using Drosophila 28S rDNA as a template, which shows the characteristic “triple-U” pattern in which the U3486 base is labelled in blue where primer extension reactions stall, and the position for the actual N^3^-modified U3485 base is labelled in green. **D. Primer extension detection of m^3^U in eukaryotic rRNAs.** At the left is a gel showing the results from primer extension using total RNA purified from different organisms with their names listed on top. At the right is a similar sequencing gel as the one in **C**. **E. The mung bean nuclease protection assay for isolating nucleotides of interest.** At the top is schematic representations of the assay, in which a DNA primer anneals with the rRNA at the position of the m^3^U base, marked with a lollipop. The primers are designed so that it covers the m^3^U base (situations illustrated at the left and the right, “m^3^U protection”), or it does not (situation in the middle, “no protection”). After nuclease digestion, gel purification of the protected RNA fragments and a further round of nuclease digestion, individual nucleotides are subjected to HPLC or Mass-Spec analyses. At the bottom is an EtBr-stained gel showing the various products after mung bean nuclease digestion. **F. RP-HPLC analyses of m^3^U.** Nucleotides recovered from the assay shown in **E** were subjected to RP-HPLC. At the left, nucleotides were purified from *ptch* heterozygotes (blue line), and homozygotes (green and black lines). A commercial m^3^U sample was spiked in the “green” sample to mark the running position of the m^3^U base. In the middle, nucleotides were purified from wildtype adults with the two primers used in the protection assay as illustrated in **E**. At the right, nucleotides were purified from yeast, mouse and human rRNA, and subjected to RP-HPLC. **G. Mass spectrometry detection of m^3^U**. Nucleotides were purified from *ptch* heterozygous, or homozygous adults. Triplicate samples were analyzed. n.d.: none detected.

To further confirm the presence and map the location of m^3^U on rRNAs from fly and human, we employed a nuclease protection principle to enrich small RNA fragments that either contain or do not contain the modified U base (Yang et al. 2016, Figure 2E), and subjected them to either HPLC or LC-Mass analyses. As shown in Figure 2F, HPLC analyses confirmed the presence of m^3^U in the *wt* RNA fragments that include U3458 but not those without. Similarly, m^3^U is present in mouse and human but not yeast rRNA, consistent with results from primer extension experiments shown in Figure 2D. Importantly, m^3^U is missing in RNA fragments containing U3458 but isolated from *ptch* mutants (Figure 2F). LC-Mass analysis also confirmed the presence of the m^3^U base in *wt* but not *ptch* mutant rRNAs (Figure 2G). Therefore, multiple methods were employed to confirm the presence of m^3^U at U3485 of fly 28S rRNA, and the presence of this modification depends on Ptch.

Ptch is homologous to the human C9orf114/SPOUT1 protein. We used RNAi to knock down its expression in HeLa cells, and used RTL-P to confirm that m^3^U on wildtype 28S rRNA impedes reverse transcription and this impediment is relieved when SPOUT1 expression is reduced (Figure S2C). Therefore, the role of Ptch/SPOUT1 in controlling m^3^U modification of 28S rRNA is evolutionarily conserved.

### Loss of m^3^U at PTC affects translational efficiency

The Ptch-regulated U3485 base is situated in a highly conserved region of PTC of the large ribosome. The Minute phenotypes carried by *ptch* mutant adults suggest defective protein synthesis (Figure 1C). To investigate the efficiency for peptide synthesis *in vivo*, we used a puromycin-based method. The antibiotics puromycin interacts with the ribosomes and gets incorporated into the elongating polypeptide chain causing its termination. The OPP derivative of puromycin, engineered with different labels, allows the detection of the newly synthesized polypeptide chains after a click chemistry reaction in both cultured cells and animal tissues (Liu et al. 2012; Lee et al. 2018).

Animals with the *ptch^1-3^* null allele die as third instar larvae. We then treated larval tissues from both the *wt* and *ptch^1-3^* animals with OPP. At different time points afterwards, a click chemistry reaction was performed to label the incorporated OPP with the Alexa488 fluor. Total proteins were extracted after labelling, and Alexa488 signals were visualized in gel, which was subsequently stained for total protein to control for equal loading. As shown in the left panels in Figure 3A, loss of Ptch reduces the amount of newly synthesized peptide. This suggests that loss of the m^3^U at PTC impairs peptide synthesis *in vivo*. However, defective peptide synthesis was not evident when we employed the OPP assay on animals mutant for the *ptch^7-7^*hypomorphic allele, regardless of whether larval tissues (Figure 3A right panels) or dissected testes and ovaries from adults (Figure 3B) were used for OPP labelling. We suspect that protein synthesis in this partial loss-of-function mutant might not be robust enough for the OPP assay to detect small decreases in translational efficiency. For example, the synthesis of specific classes of proteins rather than total protein synthesis might be disrupted by the hypomorphic mutation.

**Figure 3.**
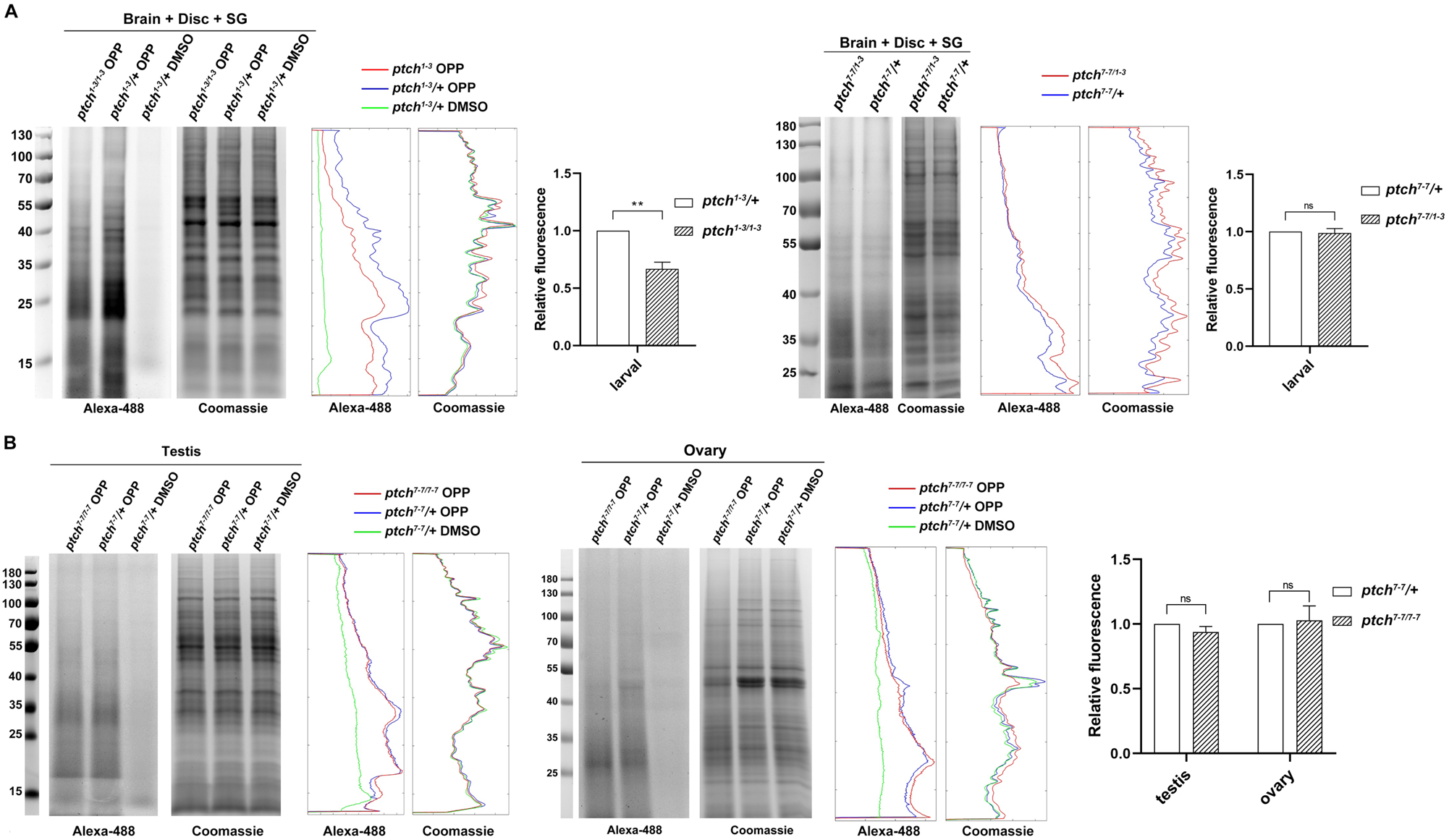
The effects of Ptch loss on translational efficiency. **A. OPP labelling of larval tissues.** The tissue samples of salivary glands (SG), imaginal discs and brains were treated with OPP for 2 hr, followed by a Click-chemistry reaction for attaching the Alexa488 fluor. Two gel pictures are shown for each experiment: the left panel shows Alexa488-OPP signals, and the right one shows the same gel stained for total protein. Markers with molecular weights in KDa are labelled. Genotypes are shown on top of the gel pictures. A negative control was included in which only the DMSO solvent of OPP was added to account for background fluorescence. Signal quantifications were included for all gel pictures. The chart to the right shown quantification of the OPP signals normalized over the total protein signals. The effects from both the null *ptch^1-3^* (left panels) and the hypomorphic *ptch^7-7/1-3^* (right panels) alleles were tested. **B. OPP labeling of adult tissues.** Testes and ovaries of the indicated genotypes were similarly treated as the larval tissues shown in **A**. Alexa488-OPP and total proteins are measured and quantified similarly as in **A**.

Mature 5.8S, 18S, and 28S rRNAs are processed from a single pre-rRNA transcript and the processing follows highly conserved and well-defined steps from yeast to human (Figure S3A, reviewed in Chan et al. 2001; Bohnsack and Bohnsack 2019). As Ptch is concentrated at the nucleolus where pre-rRNA processing happens, we determined whether this processing is abnormal in *ptch* mutants, which can be detected on Northern blots using probes from the excised rRNA fragments, such as ITS1 and ITS2. As shown in Figure S3B, processing appears normal when we used a probe from the ITS2 region. As severe processing defects could also reduce the level of mature rRNAs, we used Northern blotting (Figure S3C) and qPCR (Figure S3D) to measure all of the mature species but did not detect any change in the mutant samples. We therefore conclude that the loss of m^3^U in 28S rRNA does not significantly affect pre-rRNA processing.

### The Ptch methyltransferase is sufficient for m^3^U modification of 28S rRNA

To test whether the Ptch enzyme is solely responsible for carrying out the N^3^ modification, we set out to establish an *in vitro* methylation assay using purified components. We expressed an MBP-tagged recombinant Ptch from bacteria, and an MBP-Ptch variant that carries single amino acid mutations (Ptch^G318AG322A^ or Ptch^am^). Based on studies of other SPOUT family methyltransferases (Anantharaman et al. 2002; Michel et al. 2002; Taylor et al. 2008; Zhang et al. 2012; Hori 2017; Hernández-Cid et al. 2023), these mutations would disrupt Ptch’s interaction with its cofactor S-adenosylmethionine (SAM) and severely impede its methylase activity. The two proteins were thus purified in parallel (Figure S4A) and used in the assay.

As an RNA substrate for the reaction, we used total RNA purified from *wt* or *ptch^7-7/7-7^* mutant adults. The *in vitro* methylation reaction was followed by primer extension to detect the m^3^U base. As shown in Figure 4A, MBP-Ptch, but not MBP-Ptch^am^, is capable of installing m^3^U to mutant rRNA at the expected position, and this reaction depends on the presence of SAM. Remarkably, similarly purified human SPOUT1 recombinant protein was also able to catalyze the specific N^3^ modification of *ptch*-mutant rRNA from Drosophila (Figure 4B and Figure S4A), indicative of a strong conservation in enzyme substrate recognition. Ptch recombinant protein lacking the MBP tag (Figure S4B) also supports *in vitro* m^3^U methylation.

**Figure 4.**
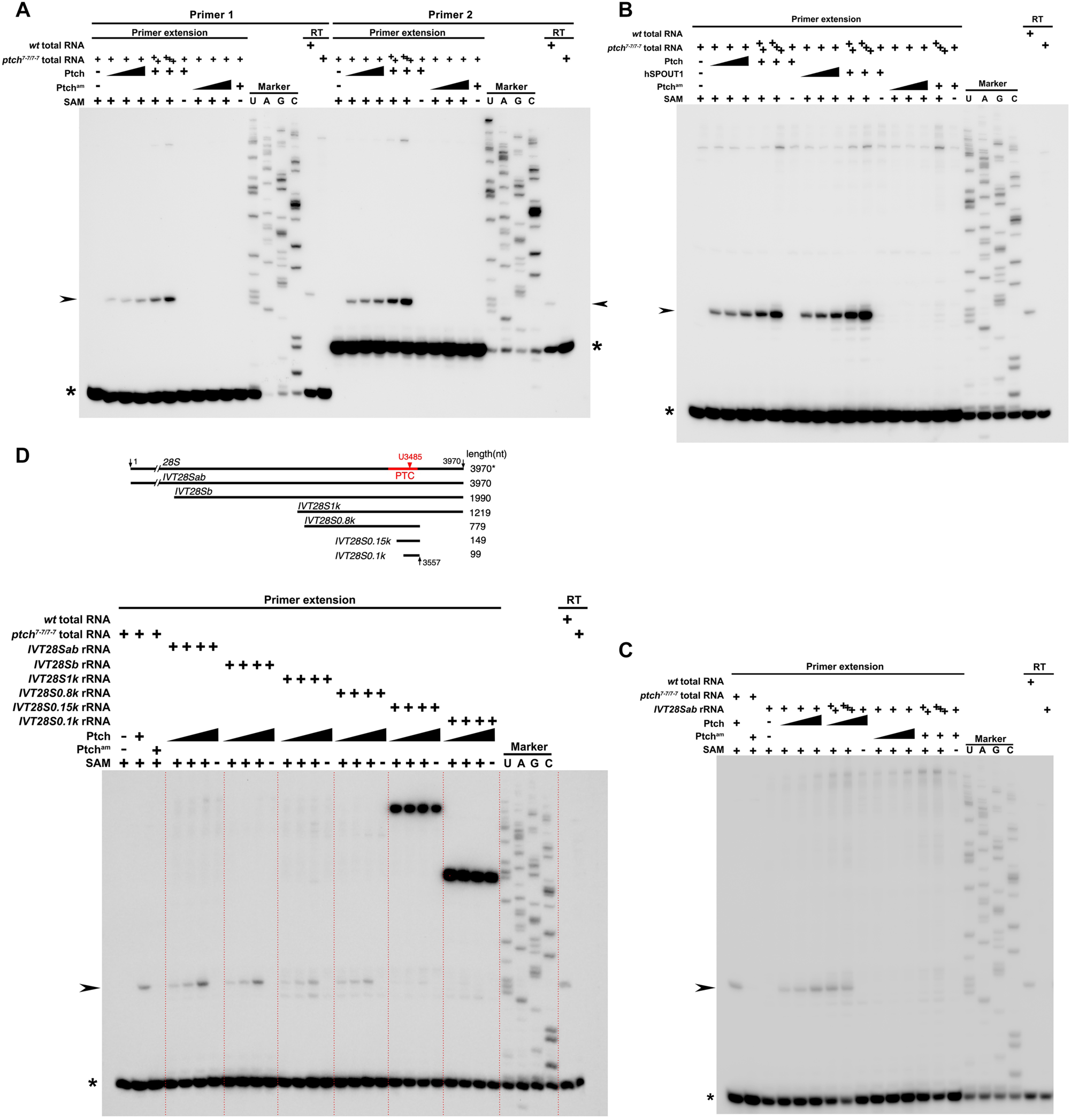
Ptch is the m^3^U methyltransferase for 28S rRNA. **A. *In vitro* methylation with total RNA.** Total RNA from *ptch* mutant adults were used in a methylation reaction with recombinant wildtype Ptch enzyme or with a mutant form of Ptch lacking a functional active site (active site mutant, Ptch^am^), both tagged with MBP. The products of the *in vitro* methylation reaction were subject to primer extension for the detection of m^3^U using two different primers, *hyt* (“primer 1”) and *2R* (“primer 2”), obtaining similar results. For each methylation reaction, 800ng (“+” lanes), or 1.6μg (“++” lanes) or 2.4μg (“+++” lanes) of total RNA were used. MBP-Ptch concentrations were 0.5, 1.0, 4.0μM (for the gradient) and 4.0μM (for “+” lanes). The MBP-Ptch^am^ concentrations were 2.0, 4.0,12μM (for the gradient) and 12μM (for “+” lanes). The SAM cofactor was used at 1.0mM (“+” lanes). “U A T C” lanes are DNA sequencing markers. The rightmost two lanes are primer extension results without the preceding methylation reaction. RT: RNA samples are directly subject to reverse transcriptase mediated primer extension without methylation reaction. Only the wildtype RNA samples yield a stalled product at the m^3^U position, marked with arrowheads. The asterisk marks the running position of the free oligos. **B. The human enzyme is functional on Drosophila substrate RNA.** *In vitro* methylation reaction was performed similarly to the one shown **A** except the following ingredients. MBP-Ptch concentrations were 30, 60, 120μM (for the gradient) and 120μM (for “+” lanes). The MBP-Ptch^am^ concentrations were 0.1, 0.2, 0.6mM (for the gradient) and 0.6mM (for “+” lanes). A recombinant human MBP-hSPOUT1 enzyme was used at a concentration of 30, 60, 120μM (for the gradient) and 120μM (for “+” lanes). SAM was used at 0.5mM (“+” lanes). Primer extension was carried out with the *hyt* primer. **C. Ptch is functional on an RNA substrate made *in vitro*.** *In vitro* methylation reaction was performed similarly to the one shown in **A** and **B** except the following ingredients. A reaction was included with the full length 28S rRNA transcribed *in vitro* (*IVT28Sab*, see **D** for details) used as the RNA substrate at the following concentrations: 0.05μM (“+” lanes), 0.15μM (“++” lanes), and 0.25μM (“+++” lanes). A control methylation reaction with *ptch* mutant RNA was included and ran at the leftmost lane. Primer extension was carried out with the *hyt* primer. **D. Preliminary investigation of substrate specificity by RNA truncation.** At the top is a schematic representation of the *in vitro* transcribed (IVT) RNA fragments used in the methylation assay shown at the bottom. The names of the RNA substrates are listed either at the top or to the left of the thick line representing each fragment drawn approximately to scale. The sizes of the fragments are listed to the right. “28S” denotes 28S rRNA purified from adults. 3970*: the size of 3970nt denotes the predicted size of the mature 28S rRNA whereas 28S from purified flies are often self-cleaved yielding a fragment approximately the same size as the IVT28Sb fragment. Key positions are labelled with arrows with the nucleotide numbers given. The approximate position of the PTC region where the m^3^U3485 lies is shown in red with the methylated base labelled with a red arrowhead. At the bottom*, in vitro* methylation reaction was performed similarly to the one shown **A** and **B** except the following ingredients. 28S fragments were used at a concentration of 0.05 μM. Primer extension was carried out with the *hyt* primer.

To further investigate how Ptch recognizes its substrate, we used *in vitro* transcribed fly 28S rRNA as substrates. In addition to the full length 28S rRNA (3970nt), we synthesized several shorter derivatives that are progressively truncated at the 5’ end (Figure 4C and 4D): a 1990nt fragment corresponding to a spontaneously generated 28S sub-fragment (Figure S3A), a 1219nt and a 779nt fragments all containing the U base of interest. Remarkably, results from the *in vitro* methylation assay largely recapitulate those from using the *in vivo* rRNA as the substrate (Figure 4C and 4D). We were encouraged by these initial positive results, and synthesized two even shorter RNA fragments centering around the modified U base (a 149nt, and a 99nt fragment). Unfortunately, neither of these two shorter fragments could be efficiently used as substrates for Ptch methylation (Figure 4D). In summary, our results suggest that the primary RNA sequence in combination of RNA structural elements are determinants of substrate recognition by Ptch. Although we have not been able to determine the minimal requirements for substrate recognition, we succeeded in reconstituting the entire m^3^U reaction with *in vitro* components paving the way for in depth enzymology studies of Ptch/SPOUT1 in the future.

## Discussion

Over thirty years have passed since it was shown that the 28S rRNA from human is N^3^ modified at a single U base. The enzyme responsible for this modification remains unknown until now. Here we identified Ptch as the responsible enzyme for the m^3^U modification on 28S rRNAs from higher eukaryotes. This modification is situated at the heart of the PTC domain of the large ribosome. Drosophila cells without this modification experience reduce protein synthesis capacity that ultimately leads to the death of the organism. Human cells deficient for the homologous enzyme do not survive as shown by others.

### m^3^U modification of PTC is evolutionarily ancient

m^3^U modification is a conserved rRNA modifications, appearing in all kingdoms of life. Bioinformatic analyses suggest that m^3^U formation is likely catalyzed by the SPOUT class of RNA methyltransferases (Anantharaman et al. 2002; Tkaczuk et al. 2007). An m^3^U methylase likely represents one of three SPOUT-class and rRNA modifying enzymes present in the Last Universal Common Ancestor (the other two being methylases for 2’-O-methylation and m^1^G). The bacterial RsmE enzyme and Bmt5 and Bmt6 enzymes from yeast catalyze m^3^U formation on rRNAs from their respective species at different positions.

At the PTC center of the large ribosome, the sequence of “5’-GGG**U**U” is universally conserved in which the modification on the first U is of interest here. The functional importance of this U residue has been extensively investigated in a myriad of mutational studies in bacteria (e.g. Porse et al. 1996; Green and Noller 1999; Youngman et al. 2004; d’Aquino et al. 2020; reviewed in Polacek and Mankin 2005). Of particular interest is the finding that 2’-O-methylation mis-targeted to this U base has physiological consequences in yeast (Liu et al. 2008). Results from our study and others before suggest that this unique m^3^U modification is present to animals, missing in yeast and possibly plants. Although it is missing in bacteria, it is present in Archaeon. A single m^3^U base has been identified in 23S rRNA of the hyperthermophile *Sulfolobus solfactaricus* although the modified residue was not identified precisely (Noon et al. 1998). A careful re-evaluation of rRNA modifications in *Haloarcula marismortui* identified an m^3^U at PTC being one of only eight chemically modified sites in its 23S rRNA (Kirpekar et al. 2005). Remarkably, the EMS-induced EL-2 mutant strain of *Halobacterium salinarium* was shown to be specifically missing the U-carried modification at PTC, which results in resistance to the antibiotics Sparsomycin (Lázaro et al. 1996). Unfortunately, neither the nature of the U-carried modification nor the responsible enzyme mutation have been identified in EL-2. Nevertheless, an enzyme responsible for this m^3^U modification has been speculated for *Haloferax volcannii* (Grosjean et al. 2008). Therefore, m^3^U modified PTC carries an ancient function that might have been replaced or made dispensable in certain species such as bacteria, yeast and plants. The identification of the m^3^U methylase here would spur renewed interests in this ancient modification.

### The function of the m^3^U modification at PTC

Although the solution conformations of 3-methyluridine have been solved (Desaulniers et al. 2005), there is a lack of studies on the effect of m^3^U on RNA chemistry. *In vivo* studies of m^3^U on bacterial and fungal rRNAs yielded limited functional insights likely due to the lack of a strong physiological consequence in the enzyme mutants. On the contrary, the loss of a single methyl group at PTC causes lethality to human cells and Drosophila adults as we showed here. From our characterization of the Drosophila mutants lacking the m^3^U modified PTC, we surmise that m^3^U is not important for the proper processing of pre-rRNA, but that normal translational proficiency measured by OPP incorporation into nascent peptides relies on its presence. However, the loss of total translational proficiency seems small.

Our last proposition is consistent with results from the characterization of the Archaean EL-2 mutant (Lázaro et al. 1996). Ribosomes purified from the EL-2 mutant had a reduced efficiency in directing translation from a polyU RNA template. Interestingly, this reduction can be compensated by increasing the concentration of Mg^2+^ in the reaction, while wildtype ribosomes loss their translational efficiency in the presence of heightened Mg^2+^ concentrations. Furthermore, the level of mature 70S ribosomes is reduced in EL-2, yet again this reduction can be counteracted with increasing Mg^2+^. Ribosomes is an important store of intracellular Mg^2+^, where it interacts with RNA and proteins, and is important for the overall structure of ribosome (e.g., Misra and Draper 1998; Schneider et al. 2012; Nierhaus 2014). Therefore, the loss of m^3^U at PTC might have altered a specific structure of ribosome. This conclusion is further supported by the Archaean study in which the EL-2 mutation also changes the chemical accessibility of the neighboring U base, another universally conserved base with demonstrated importance. The combination of facile genetics in archaeal models and the recently developed system for introducing rRNA variants *in vivo* (Jüttner et al. 2020), are anticipated to facilitate the molecular characterization of m^3^U’s effect on ribosomal structure and function.

### Substrate recognition by Ptch

Another significant contribution from our study is the demonstration that the Ptch enzyme is both necessary and sufficient for catalyzing the specific m^3^U modification. In particular, we succeeded in reconstituting the methylation reaction with recombinant components. In contrast, only partially purified ribosomes can serve as substrates for m^3^U methylases from bacteria or yeast (Basturea and Deutscher 2007; Sharma et al. 2014). Remarkably, recombinant human enzyme is capable of using the Drosophila RNA as a substrate indicating that the determinants for substrate recognition must be conserved.

Although *in vitro* synthesized 28S rRNA fragments longer than 700nt serve effectively as substrates for Ptch, significantly shorter RNA molecules have not produced positive results suggesting that RNA sequence alone is not sufficient for substrate recognition. Whether secondary structure(s) constitutes an essential element for such recognition requires further investigation.

### Ptch as a potential target for “cancer antibiotics”

Most of the natural occurring antibiotics target the ribosome. Remarkably, PTC is by far the most targeted region (Polacek and Mankin 2005), suggesting PTC is a naturally selected target for growth inhibition. Our study identified a fine tuner for ribosomal function in which the presence or absence of a single methyl group dictates cell survival in complex organisms. A structural and chemical analysis of a large number of human RNA methyltransferases led to the conclusion that Ptch/C9orf114 is one of the most “druggable” enzymes for having a cofactor binding pocket with the least charges and the highest extent of enclosure (Schapira 2016). Proliferation of cancer cells demands heightened ribosomal functions, and the nucleolus has been championed as an emerging target for cancer therapy (Hein et al. 2013). We speculate that small inhibitors of Ptch would deliver a highly focused disruption of ribosomal function potentially making their toxicity to healthy cells more manageable.

## Materials and Methods

### Fly stocks, yeast strains and animal cells

The following stocks were obtained from the Drosophila Bloomington Stock Center at Indiana, USA: BL#52669 has a *vasa*-driven *cas9* transgene producing Cas9 protein in the germlines; BL#9539 (*Df(2R)BSC152/CyO*) carries a chromosomal deficiency of the *ptch* region. The *w^1118^*stock was used as wildtype. The construction of *ptch* mutant lines, and transgenic lines carrying *ptch* rescuing constructs are described below and illustrated in Figure 1A. Drosophila stocks and crosses were maintained on a standard cornmeal media and kept at 25°C.

The following samples were used for m^3^U detection of in rRNA from other species: SK1 strain of *Saccharomyces cerevisiae*; h90 strain of *Schizosaccharomyces pombe*; blood from mouse C57BL/6 strain; and HeLa human cells.

### CRISPR/Cas9-mediated mutagenesis of ptch

Cas9 induced *ptch* mutations were obtained using a transgenic approach in which Cas9 and sgRNA were expressed from two transgenes carrying either the *vasa* promoter for Cas9 expression or the *U6* promoter for sgRNA expression. sgRNA#1 with the sequence of 5’-ATTGCTTTAGCGGCTCTCC**AGG** (PAM sequence in bold), was used to generate frameshift mutations of *ptch^7-7^* (an insertion of 8bp accompanied by a deletion of 4bp, potentially encoding a truncated Ptch missing the last 116 residues), and *ptch^12-2^* (a 2bp deletion, potentially encoding a C-terminally truncated Ptch protein). sgRNA#1 together with sgRNA#2, with the sequence of 5’-GGGAACAGCTATGCTCAGCG**TGG** (PAM in bold), were used to generate the deletion mutation of *ptch^1-3^*(a 955bp deletion). All mutations were verified by PCR amplification followed by sequencing using DNA from homozygous animals as templates. PCR primers used for mutation detection are listed in Table S1.

### Construction of rescuing transgenic lines of *ptch*

A 5.8 kb DNA fragment from the genomic region of *ptch* (nt 10025839-10031659 of chromosome *2R* in release dm6) that contains the endogenous regulatory elements of *ptch* was cloned into the vector of pUASTattB. Starting from this rescuing construct, we generated the *ptch^G318AG322A^* active site mutation by site-directed mutagenesis. Also starting from this construct, we inserted an *egfp* gene to the end of *ptch* before the STOP codon by bacterial recombineering, resulting in a transgene making GFP tagged Ptch proteins. All transgenic constructs were inserted at 86F of chromosome *3R* by phiC31 integrase-mediated germline transformation, and were subsequently introduced into a *ptch* mutant background by genetic crossing. PCR primers for plasmid construction and verification are listed in Table S1.

### Fluorescence in Situ Hybridization (FISH) and immunostaining

Salivary glands, ovaries and testes of *ptch^gfp^* flies (full genotype: *ptch^1-3^/^1-3^; [ptch^gfp^]/[ptch^gfp^]*) were dissected in PBS. Tissues were fixed in 4% paraformaldehyde for 4hr. After washed three times with 2X SSCT (0.1% Triton X-100), they were dehydrated using an ethanol gradient. After air drying, pre-hybridization was performed in 25% formamide, 5X Denhardt, 5% dextran sulfate, 2X SSC, 0.2% BSA, and 125μg of tRNA for 2 h at 37°C. Tissues were incubated at 37°C overnight with a final probe concentration of 0.5 ng/mL. Post-hybridization washes were performed three times in 0.2X SSCT for 15 min each. Tissues were then stained with DAPI (0.2 μg/mL in 4X SSCT) for 5 min, washed briefly in 4X SSCT, and allowed to air dry. Fluorescent images were captured by an Olympus IX83 confocal microscope. Probes are listed in Table S1.

Salivary glands, ovaries and testes of *w^1118^* and *ptch^gfp^* were dissected in cold PBS. Tissues were fixed with a mixture of 3.7% formaldehyde diluted in PBS for 15 min. Heptane was subsequently added as a treatment for 30 min at room temperature. After washing three times with 1X PBST (0.1% Triton X-100), a mouse anti-Fib antibody (from Abcam) was incubated overnight at 4°C. After washing three times with 1X PBST (0.1% Triton X-100), Alexa Fluor 555 conjugated goat anti-mouse IgG (1:200, Abcam) and DAPI were incubated for 1 h. Fluorescent images (GFP fluorescence and anti-Fib signals) were photographed by an Olympus IX83 confocal microscope.

### Analysis of rRNA by Northern blotting and qPCR

Northern hybridization analysis of 18S and 28S rRNAs was performed with a protocol based on one described by Sharma et al. (2014). Total RNA was prepared by TRIzol reagent (Invitrogen, Carlsbad, California CA, USA). Ten μg of total RNA each from adults of *w^1118^*, *ptch^7-7/12-2^* and *ptch^7-7/12-2^*; *[ptch^+^]* were separated on a 1% agarose gel in 1X TAE supplemented with 6.66% formaldehyde and transferred to a positively charged nylon membrane (Hybond N+, GE Healthcare) using capillary transfer. Northern hybridization analysis of 5.8S and 5S rRNAs was performed with a protocol based on one described by Brown et al. (2004). Agarose (1g) was dissolved in 72mL of water and cooled to 60°C in a water bath, followed by the addition of 10mL of 10X MOPS running buffer and 18mL of 12.3 M formaldehyde. Ten μg of total RNA from adults were dissolved in 11μl of water, and mixed with 5μl of 10X MOPS running buffer, 9μl of 12.3 M formaldehyde, 25 μl formamide, and 10μl of formaldehyde loading buffer. The mixture was briefly centrifuged to collect the liquid and incubated for 15 min at 55°C. After running the gel, it was placed in an RNase-free glass dish and soaked for 45 min in 10 gel volumes of 20X SSC buffer. The RNA was transferred from the gel to a nylon membrane overnight. The membrane was rinsed in 2X SSC and allowed to dry, followed by immobilization of RNA with a UV transilluminator. The membrane was incubated with biotin-labeled DNA probes for 3 h at 42°C with rotation, washed three times with 2X SSC at room temperature, after which 0.1% of sodium dodecyl sulfate (SDS) was added. A chemiluminescent biotin-labeled nucleic acid detection kit (Beyotime, China) was used to detect the signal using a manufacture provided protocol.

Total RNA from five female adults of *w^1118^*, *ptch^7-7/12-2^*, and *ptch^7-7/12-2^*; *[ptch^+^]* were extracted using TRIzol reagent. cDNA was generated from 1μg of total RNA using the PrimeScriptTM RT reagent kit (TaKaRa, Shiga, Japan) for the quantitation of rRNA by quantitative real-time PCR. A Bio-Rad CFX96 Real-Time System (Bio-Rad, Hercules, CA, USA) and a KAPA SYBR FAST Universal qPCR Kit (Kapa Biosystems, Boston, MA, USA) were used. The cDNA standard for each gene was diluted to different concentrations and used as templates for PCR reactions. The expression of *rp49* was used as a control. The standard curves of these genes were drawn with the logarithm of the copy number of template DNA as the abscissa and the measured CT value as the ordinate. The initial copy number of 18S rRNA, 28S rRNA, 5.8S rRNA, or 5S rRNA was calculated from the CT value of each cDNA sample after adjustment with *rp49*. Each quantification was done in three biological replicates. The primers are listed in Table S1.

### Primer extension for the detection of m^3^U

Two 5’ biotin-labelled DNA primers (HYT and 2R) were used for primer extension. A primer/RNA pre-mixture containing 8μl of RNase-free water, 1μl of primer (one picomole) and 1μg of template RNA per reaction was denatured at 70°C for 3 min then chilled on ice. After incubation at 42°C for 10 min, 5X M-MLV reverse transcription buffer, dNTP mix (final concentration of 1 mM), 200 U of M-MLV reverse transcriptase (Promega) and RNase-free water were added to make up a 25μl reaction. The primer extension reaction was performed at 42°C for 1hr, and DNA products were separated on 40% polyacrylamide-8M urea gels. To generate markers for the primer extension products, a DNA sequencing reaction was performed by a standard protocol. Briefly, the 28S ribosomal DNA of *Drosophila melanogaster* was PCR amplified and cloned into the pMD-18T vector. A 15μl sequencing reaction mixture contains 1μg of the above plasmid, 10 U of Taq DNA polymerase (Promega), 5μl of a ddNTP mixture, 3μl of 5X Taq buffer, and 1 pmol of the biotin-labeled primer. This marker was run along with the primer extension products, transferred to nylon membranes and developed using streptavidin-HRP.

### Small RNA fragments purification by mung bean nuclease protection

A mung bean nuclease protection assay was modified from a protocol described in Yang et al. (2016). Briefly, a 175μl of DNA/RNA mixture, containing 10μl of the protecting DNA oligonucleotides (DNA oligos) and 100μg of total Drosophila RNA, was incubated with 10% DMSO in 0.3 volume of hybridization buffer (250 mM HEPES, 500 mM KCl, 5% DMSO at pH 7) at 90°C for 5 min, then slowly cooled to room temperature over 2 h. After this hybridization step, 20μl of 10X mung bean nuclease buffer (New England Bio Labs, NEB) and 4μl of the nuclease (NEB) along with 1μl of RNase A (Sigma) were added to the digestion and allowed to react for 1hr at 35°C. After the digestion step, the protected fragment (DNA-RNA hybrid) was extracted by phenol/chloroform extraction, followed by ethanol precipitation overnight. The protected rRNA fragments were separated from the complementary DNA oligos on a denaturing 13% polyacrylamide-8M urea gel, and were excised and eluted in elution buffer (1 mM ethylenediaminetetraacetic acid, 0.1% sodium dodecyl sulphate and 0.4 M NaOAc, pH 5.2) at 25°C for 4 h. 3 volume of ethanol was added to precipitate RNA fragments. The sequences of the protecting DNA oligos were listed in Table S1.

### RTL-P, RP-HPLC, and Mass spectrometry analyses of m^3^U

Reverse transcription at low deoxyribonucleoside triphosphate (dNTP) concentrations followed by polymerase chain reaction (RTL-P) was prepared as described by Dong et al. (2012). The specific RT primer and two forward primers for PCR amplification were listed in Table S1.

For Reversed phase high performance liquid chromatography (RP-HPLC) analysis of m^3^U modifications, the protected RNA fragments from the mung bean nuclease protection assay were denatured for 2 min, followed by rapid cooling on ice. After digestion to completion by nuclease P1 and alkaline phosphatase, the hydrolysate was analyzed by HPLC according to the method described in Yang et al. (2016). Briefly, nucleosides were loaded on a Supelcosil LC-18-S HPLC column (25 cm X 4.6 mm, 5 μm) at 25°C on a Dionex UltiMate 3000 HPLC system (Thermo Fisher Scientific). For m^3^U detection, the 254 nm of absorption peak wavelength was used. The elution conditions for m^3^U were changed to an gradient mode using 100% buffer A (10mM of NH4H2PO4, 2.5% of methanol, pH 5.3) for 20 min, 90% buffer and 10% buffer B (10mM of NH4H2PO4, 20% of methanol, pH 5.1) for 5 min, 75% buffer A and 25% buffer B for 11 min, 50% buffer A and 50% buffer B for 9 min, and 100% buffer B for 20 min. Buffer C (10mM of NH4H2PO4, 35% of acetonitrile, pH 4.9) was used to clean and recycle the column after each run. The flow rate was 1 mL/min.

Mass-spec analyses were performed by the Wuhan Greensword Creation Technology Co. Ltd., (Wuhan, China) based on UHPLC-MS/MS analysis (Thermo Scientific Ultimate 3000 UHPLC coupled with TSQ Quantiva). In brief, the protected RNA fragments were enzymatically digested according to a previously described method (Shen et al., 2015), and subsequently was subjected to UHPLC-MS/MS analysis.

### Knockdown of human SPOUT1 by RNA interference

siRNAs were obtained from RIBBIO (Guangzhou, China). HeLa cells were plated on 24-well plates and grown to 50% confluence. Cells were transfected using Lipofectamine RNAiMAX (Invitrogen) with 100 nM of NC siRNA or SPOUT1 siRNAs (si-1: GCAGGACCCUCGCACCAAA; si-2: GCACCAGGAUCUACAGUUU; si-3: GUGUCCUCUUUGACCUGUA). After 48, 72, 96 h of growth, cells were lysed with Trizol. Total RNA was reverse transcribed to detect the expression level of *SPOUT1* with *gapdh* used as a control. Additionally, total RNA was reverse transcribed with specific primers for the detection of m^3^U by RTL-P. Three biological replicates were included. The primers are listed in Table S1.

### Op-puro labeling of cells and protein detection by fluorescence

Labelling was performed with Click-iT® Plus OPP Protein Synthesis Assay Kits (Thermo, C10456) and Click-iT® Protein Reaction Buffer Kit (Thermo, C10276) on tissues. Tissues (brain, disc and salivary gland of larvae, testis and ovary of adult flies) were dissected in Schneider’s medium, transferred to Schneider’s medium with 20 μM OPP, and incubated for 0.5 or 2 hr. Samples were washed with PBS briefly and then homogenized in lysis buffer (1% SDS, 50 mM Tris-HCl, 1 mM MgCl2, 1μl Benzonase, protease inhibitor cocktail). After 5 min of lysis, a click reaction was performed per the protocol from kit (Thermo, C10276) for 30min, with continuously vortexing. Following centrifugation for 3 min, the supernatant was transferred to a new tube with Laemmli buffer and heated for 10min at 98℃. Protein samples prepared as above were subjected to SDS-PAGE electrophoresis. The gel was fixed with a solution of 60% methanol and 30% acetic acid, detected for Alexa-488 and subsequently stained with Coomassie blue for total protein quantification. Experiments on different genotypes were performed in at least three biological replicates. Different durations of OPP incubation (0.5 or 2 hr) yield similar conclusions.

### Expression and purification of Ptch recombinant proteins

The ORF sequences of Ptch and Human Spout1 were cloned into the expression vector pLOU3 (Qi et al. 2018), in which the target protein is tagged at its N terminal with a His6 tag followed by an MPB tag. Site-directed mutagenesis was employed to introduce the *ptch^G318AG322A^*active site mutation into the above pLOU3-Ptch vector. The expression plasmids were transformed into *E. coli* BL21(DE3) for IPTG (0.5 mM) induced protein expression. Soluble proteins were purified by a standard Ni-NTA resin-based method and eluted with imidazole. Eluted proteins were buffer changed against PBS with 600mM Trehalose and 30% glycerol at RT through PD SpinTrap G-25 columns (Cytiva), and frozen in small aliquots and stored at −80°C.

### *In vitro* m^3^U methylation assay

Total RNA (800μg) from *ptch^7-7/7-7^* adults was reconstituted in 1X methylation buffer (50 mM Tris-HCl, 50 mM NaCl, 10 mM MgCl_2_, 10 mM DTT, 0.5 U of RNase Inhibitor, pH 8.0) at room temperature for 5 min. Various amount of recombinant MBP-Ptch proteins (see legends of Figure 4) was added to the RNA mixture with a final volume of 20μl, followed by the addition of 5μl of SAM in a final concentration of 0.5 mM. After incubation at 25°C for 1 h, RNA was purified from the methylation mixture by phenol/chloroform extraction, followed by ethanol precipitation overnight. Purified RNA was subject to m^3^U detection by primer extension as described above.

To synthesize RNA substrates for the methylation assay, fragments of 28S rDNA was amplified by PCR using forward primers containing promoter sequence for the T7 RNA polymerase (Table S1). 28S rRNA fragments were synthesized using a T7 RiboMAXTM Express Large Scale RNA Production System (Promega, Wisconsin, Madison, USA) following the manufacturer’s instruction. Synthesized RNA was purified by phenol/chloroform extraction followed by ethanol precipitation, and subsequently used for *in vitro* methylation as described above.

## Acknowledgments

We thank Mr. Xionghong Tan at the University of South China for his technical assistance.

## Funding

This work was supported by the National Natural Science Foundation of China (3221101328) to YSR and (32300524) to YH; the Foundation of Guangdong Academy of Agricultural Sciences (R2023PY-JX010) to JC; the National Natural Science Foundation of Hunan Province China (2023JJ40536) to MC, and a startup fund from the University of South China to YSR.

## Author Contribution

Conceptualization: JC, YB, YH, CM, YSR Methodology: JC, YB, YH, CM, YSR

Investigation: JC, YB, YH, CM, YW, ZG, XW, YL, YSR

Visualization: JC, YB, YH, CM, YSR

Supervision: YSR

Writing—original draft: JC, YSR

Writing—review & editing: JC, YB, YH, CM, YSR

## Competing Interest

All authors declare they have no competing interests.

## Data and Materials Availability

All data needed to evaluate the conclusions in the paper are present in the paper and/or the Supplementary Materials. Drosophila stocks, plasmids and antibodies are freely available upon request to YSR.

## Supplemental figure legends

**Figure S1.**
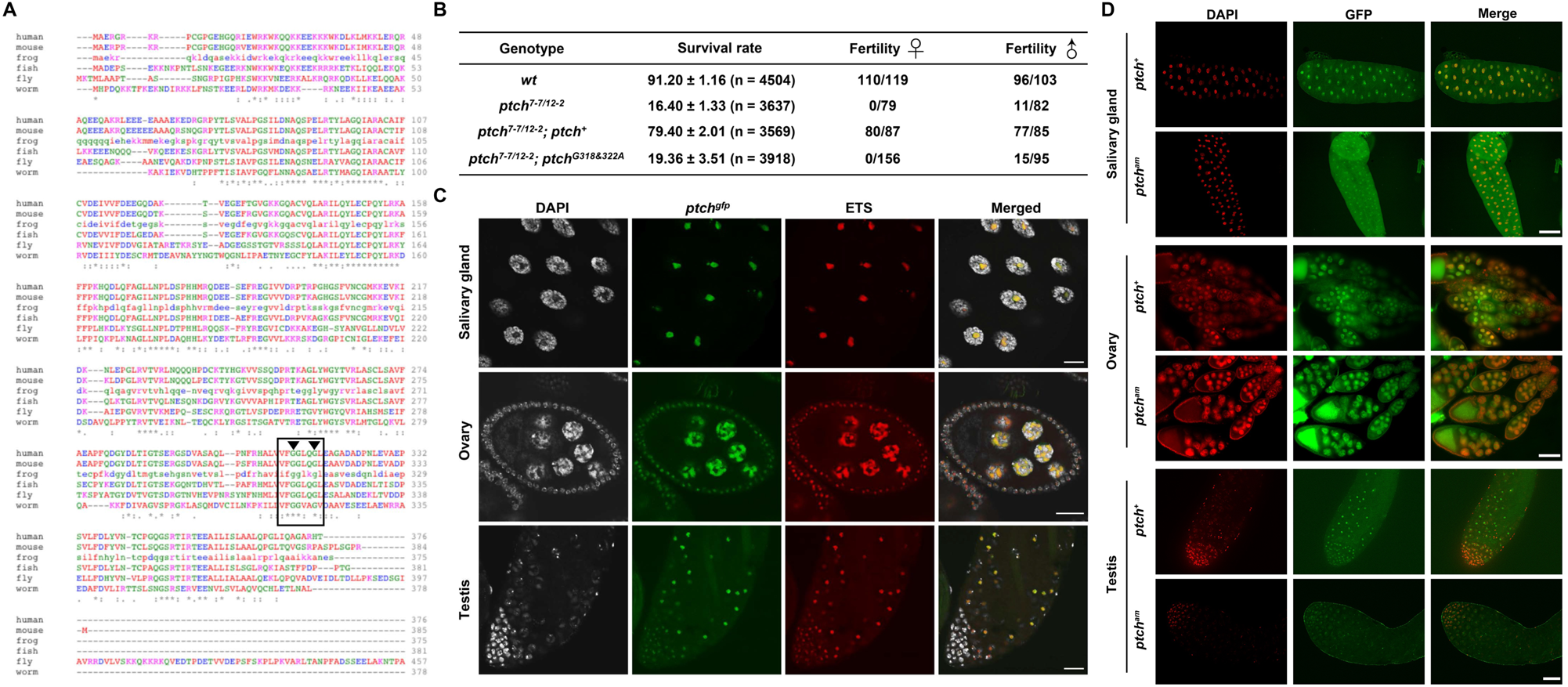
Ptch homologous proteins and Ptch’s localization in Drosophila tissues. **A. An alignment of representative Ptch homologs.** A protein alignment was generated by Clustal using the following protein sequences in the Uni-Prot database: human SPOUT1/C9orf114 (Q5T280); mouse SPOUT1 (Q3UHX9); frog SPOUT1 (Q6NRR6); fish C9orf114/cb90 (Q66ID3); fly Ptch/CG12128 (A1Z830); and worm B0361.1/Spot-1 (Q10950). The conserved region of SAM interaction has been outlined in black with the two Glycine residues marked with arrowheads, which were mutated to create the *ptch^am^* mutation. **B. Quantifications of the semi-lethal and sterility phenotypes.** Survival rates were measured in three biological replicates by crossing *ptch* heterozygous parents and scoring the homozygous survivors and the total progeny. Fertility is calculated by dividing the number of fertile adults over the number of total adults tested. **C. Ptch co-localizes with pre-rRNA in the nucleolus.** FISH experiments were conducted on dissected tissues from *ptch^gfp^* animals (complete genotype: *ptch^1-3/1-3^; [ptch^gfp/gfp^]*), using a probe to the ETS region of the pre-rRNA. In the images, DNA signals are in white, GFP signals in green, and ETS-probe signals in red. Scale bars represents 40μm. **D. The active site mutant form of Ptch localizes normally.** GFP fluorescence was used to monitor Ptch^am^ localization in various tissues. The complete genotype is *ptch^7-7/7-7^; [ptch^am/am^]*. Animals carrying the wildtype, egfp-tagged, *ptch^+^* transgene, were used for comparison. DAPI signals are in red, GFP in green. The scale bar for Salivary gland represents 200μm. The scale bar for Ovary represents 100μm. The scale bar for Testis represents 40μm.

**Figure S2.**
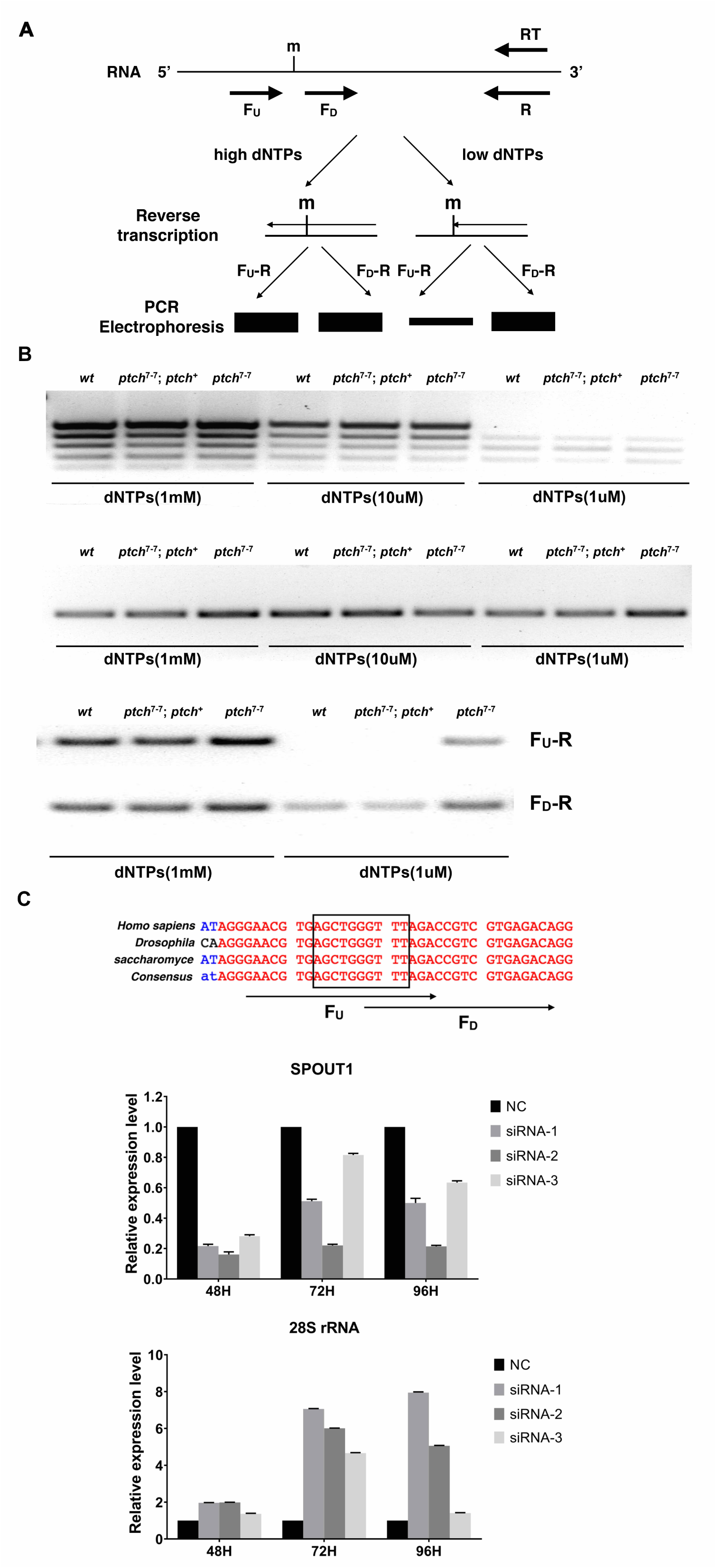
m^3^U4500 in human rRNA is controlled by SPOUT1. **A. The principle of the RTL-P method.** F_U_ denotes the primer upstream of the RNA modification (m), F_D_ denotes the primer downstream of the modification, R is the common primer, and RT is the primer used to prime reverse transcription. A restrictive amount dNTPs (“low dNTPs”), but not “high dNTPs”, impedes reverse transcription at the site of the modification. The subsequent PCR amplification would be less productive for the “low dNTPs” RT condition and using the F_U_-R PCR primer pair but not the F_D_-R pair. **B. Representative results from the RTL-P test.** At the top, a combination of five F_U_-R primer pairs assaying different regions of rRNA were used in RTL-P. Regions with modified bases can be identified but none involves the regulation of Ptch. Genotypes were listed on top of the gel picture. In the middle, an example is given in which no modification in a particular rRNA region can be detected by the RTL-P method using F_U_-R. At the bottom, a potential modification at the region of U3485 was detectable by RTL-P, and is under the control of Ptch. **C. RTL-P detection of m^3^U4500 in human rRNA.** At the top is an alignment of the region from yeast, fly and human rRNAs, and the primer design for RTL-P, with the m^3^U region highlighted in black. In the middle, qPCR was used to detect *spout1* level in RNAi experiments. At the bottom, RTL-P results show an increase of the F_U_-R products over the F_D_-R control in RNAi samples. Note that the level of RT-PCR amplification is generally anti-correlated with the extent of *spout1* knockdown.

**Figure S3.**
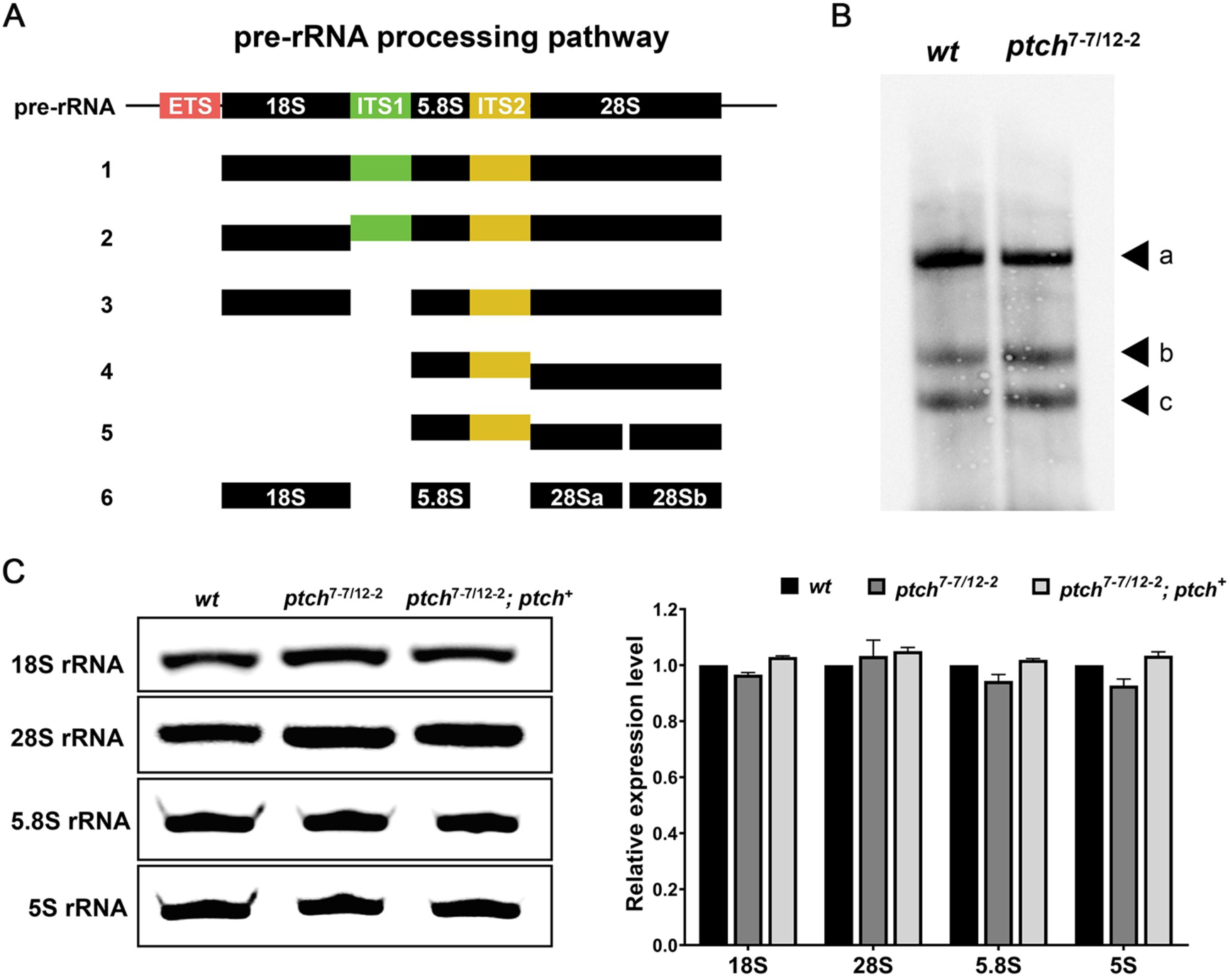
Loss of Ptch does not affect pre-rRNA processing. **A. A diagram for the steps of pre-rRNA processing in Drosophila.** The mature fragments are labelled in black, and excised ones colored. **B. pre-rRNA processing of ITS2 is normal.** A Northern blot hybridized with an ITS2 probe shows similar amounts of rRNA fragments containing ITS2 from adult total RNA. Based on the approximate molecular weights, fragment “a” corresponds to pre-rRNA, fragment “b” corresponds to ITS1+5.8S+ITS2+28S, and fragment “c” corresponds to 5.8S+ITS2+28S. **C. The level of mature rRNA is not affected by the loss of Ptch.** At the left, Northern blot was used to detect mature rRNA species in wildtype, *ptch*-mutant and *ptch*-rescued adults. At the right, qPCR was employed for the detection of rRNA fragments in adults.

**Figure S4.**
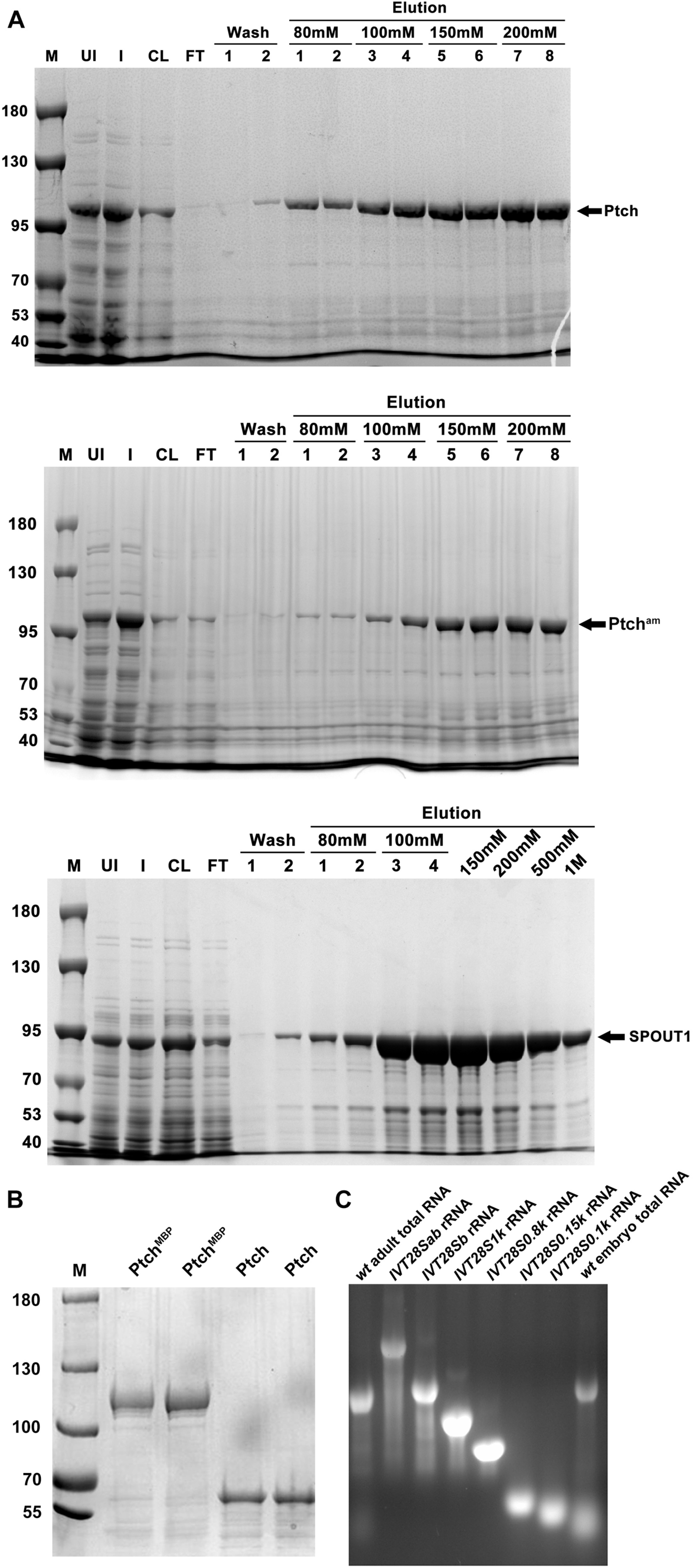
Preparations for the *in vitro* methylation assay. **A.** Purification of MBP-tagged recombinant proteins. SDS-PAGE gels stained with Coomassie Blue show the results from bacterial expression and Nickel-resin based purification of His6 and MBP-tagged Drosophila Ptch protein (top), Drosophila Ptch active site mutant protein (middle) and Human SPOUT1 protein (bottom). UI: uninduced bacterial total extract; I: total extracts induced with IPTG; CL: cleared lysate from induced extracts; FT: flow through lysate after nickel column purification. Target proteins were eluted with a buffer containing increasing amount of imidazole in mM. **B**. Successful Removal of the MBP tag. A TEV protease cleavage site between MBP and Ptch was utilized for the removal of the MBP tag. Results from two independent experiments were shown. **C**. *In vitro* transcribed rRNA fragments were run on an agarose gel and stained with EtBr. Total RNA from wildtype adults and embryos were also loaded as controls.

**Table S1.**
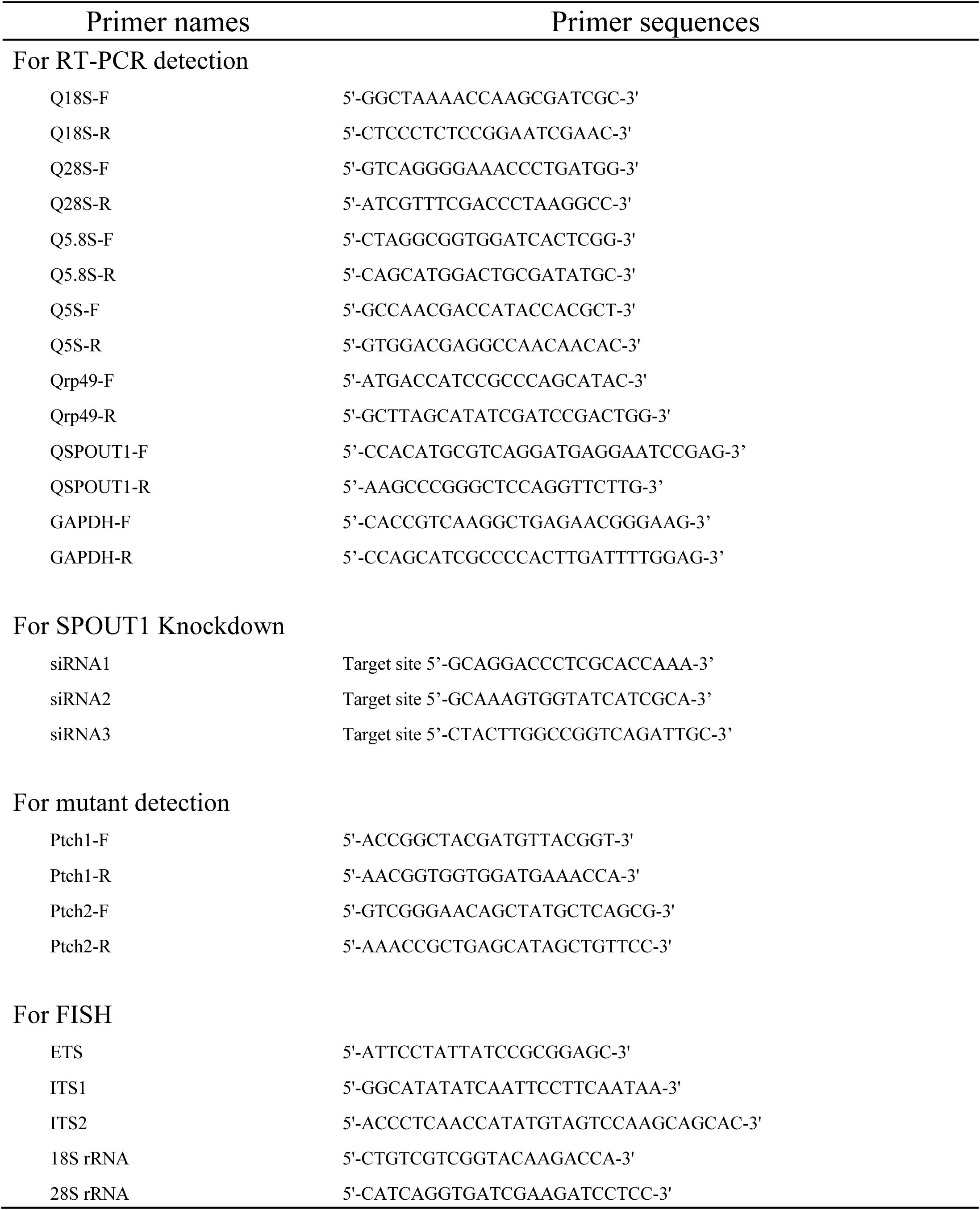
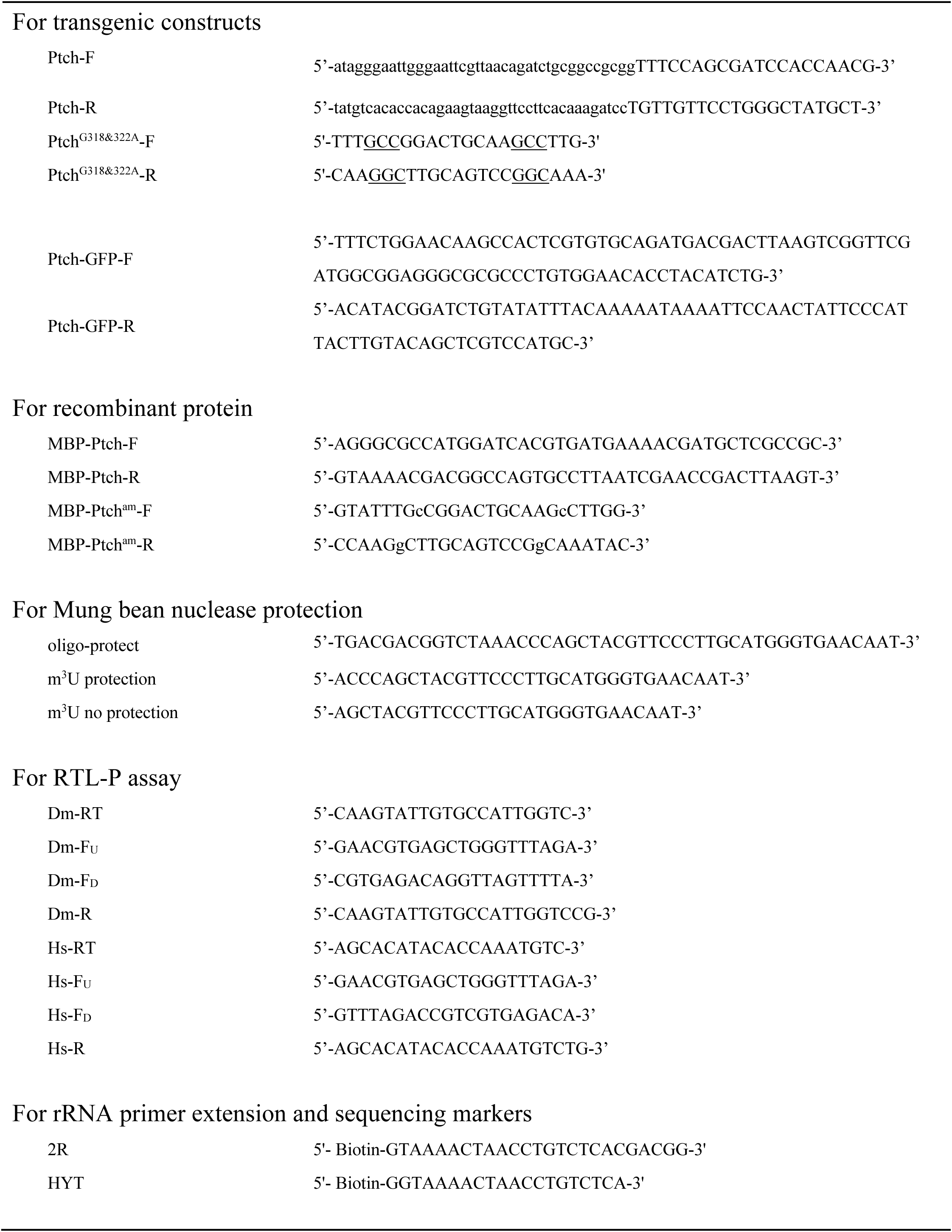
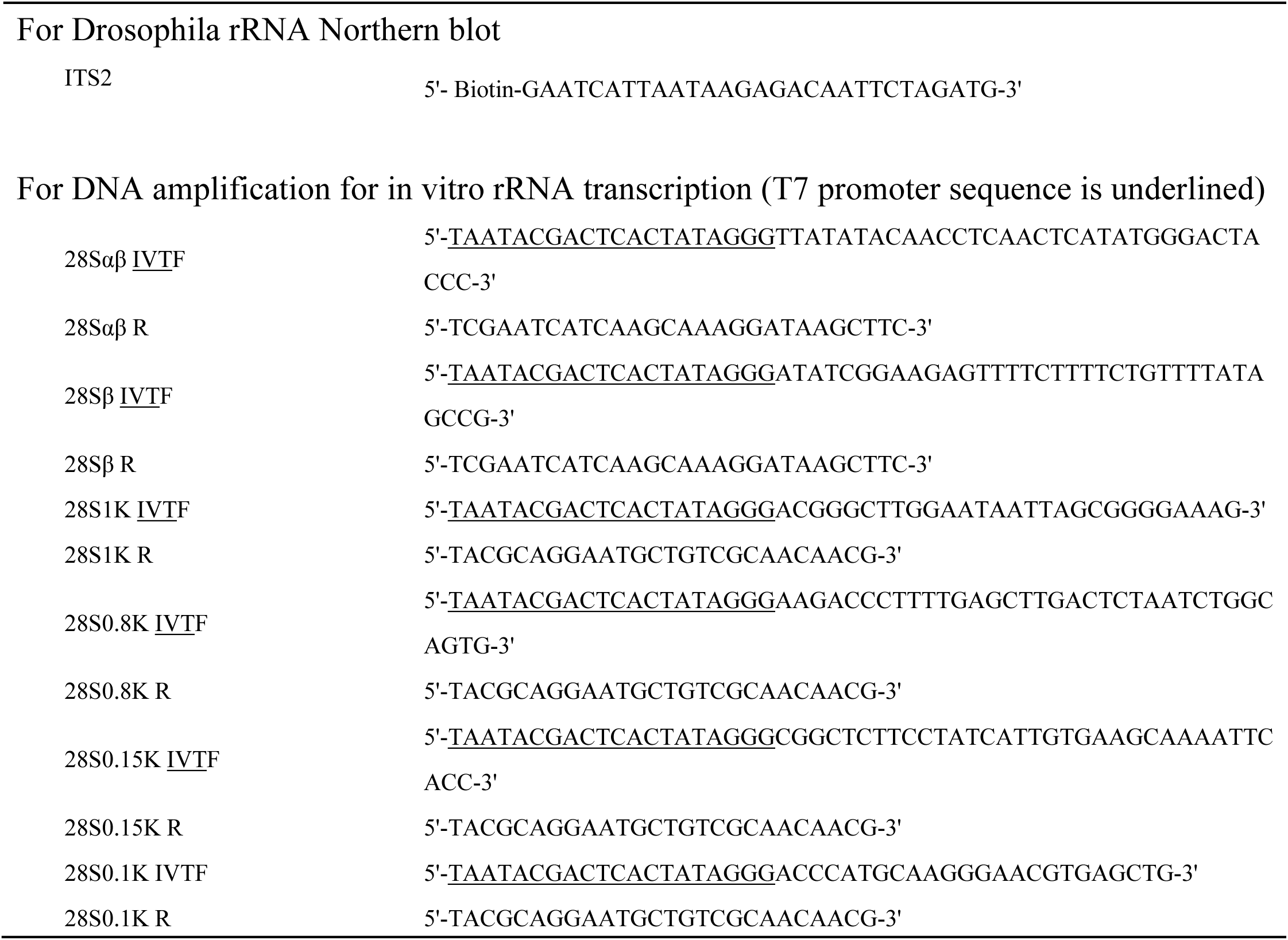
PCR primers used in this study.

## Notes

### Competing Interest Statement

The authors have declared no competing interest.

## References

Anantharaman V, Koonin EV, Aravind L. SPOUT: a class of methyltransferases that includes spoU and trmD RNA methylase superfamilies, and novel superfamilies of predicted prokaryotic RNA methylases. J Mol Microbiol Biotechnol. 2002 Jan;4(1):71–5.

Basturea GN, Rudd KE, Deutscher MP. Identification and characterization of RsmE, the founding member of a new RNA base methyltransferase family. RNA. 2006 Mar;12(3):426–34. doi: 10.1261/rna.2283106.

Basturea GN, Deutscher MP. Substrate specificity and properties of the Escherichia coli 16S rRNA methyltransferase, RsmE. RNA. 2007 Nov;13(11):1969–76. doi: 10.1261/rna.700507.

Baudin-Baillieu A, Fabret C, Liang XH, Piekna-Przybylska D, Fournier MJ, Rousset JP. Nucleotide modifications in three functionally important regions of the Saccharomyces cerevisiae ribosome affect translation accuracy. Nucleic Acids Res. 2009 Dec;37(22):7665–77. doi: 10.1093/nar/gkp816.

Baxter-Roshek JL, Petrov AN, Dinman JD. Optimization of ribosome structure and function by rRNA base modification. PLoS One. 2007 Jan 24;2(1):e174. doi: 10.1371/journal.pone.0000174.

Bohnsack KE, Bohnsack MT. Uncovering the assembly pathway of human ribosomes and its emerging links to disease. EMBO J. 2019 Jul 1;38(13):e100278. doi: 10.15252/embj.2018100278.

Bonnerot C, Pintard L, Lutfalla G. Functional redundancy of Spb1p and a snR52-dependent mechanism for the 2’-O-ribose methylation of a conserved rRNA position in yeast. Mol Cell. 2003 Nov;12(5):1309–15. doi: 10.1016/s1097-2765(03)00435-0.

Bourgeois G, Ney M, Gaspar I, Aigueperse C, Schaefer M, Kellner S, Helm M, Motorin Y. Eukaryotic rRNA Modification by Yeast 5-Methylcytosine-Methyltransferases and Human Proliferation-Associated Antigen p120. PLoS One. 2015 Jul 21;10(7):e0133321. doi: 10.1371/journal.pone.0133321.

Brown, T, Mackey, K and Du, T (2004) Analysis of RNA by northern and slot blot hybridization. Curr Protoc Mol Biol Chapter 4: Unit 4 9.

Chan HY, Brogna S, O’Kane CJ. Dribble, the Drosophila KRR1p homologue, is involved in rRNA processing. Mol Biol Cell. 2001 May;12(5):1409–19. doi: 10.1091/mbc.12.5.1409.

d’Aquino AE, Azim T, Aleksashin NA, Hockenberry AJ, Krüger A, Jewett MC. Mutational characterization and mapping of the 70S ribosome active site. Nucleic Acids Res. 2020 Mar 18;48(5):2777–2789. doi: 10.1093/nar/gkaa001.

Desaulniers JP, Chui HM, Chow CS. Solution conformations of two naturally occurring RNA nucleosides: 3-methyluridine and 3-methylpseudouridine. Bioorg Med Chem. 2005 Dec 15;13(24):6777–81. doi: 10.1016/j.bmc.2005.07.061.

Dong ZW, Shao P, Diao LT, Zhou H, Yu CH, Qu LH. RTL-P: a sensitive approach for detecting sites of 2’-O-methylation in RNA molecules. Nucleic Acids Res. 2012 Nov 1;40(20):e157. doi: 10.1093/nar/gks698.

Erales J, Marchand V, Panthu B, Gillot S, Belin S, Ghayad SE, Garcia M, Laforêts F, Marcel V, Baudin-Baillieu A, Bertin P, Couté Y, Adrait A, Meyer M, Therizols G, Yusupov M, Namy O, Ohlmann T, Motorin Y, Catez F, Diaz JJ. Evidence for rRNA 2’-O-methylation plasticity: Control of intrinsic translational capabilities of human ribosomes. Proc Natl Acad Sci U S A. 2017 Dec 5;114(49):12934–12939. doi: 10.1073/pnas.1707674114.

Green R, Noller HF. Reconstitution of functional 50S ribosomes from in vitro transcripts of Bacillus stearothermophilus 23S rRNA. Biochemistry. 1999 Feb 9;38(6):1772–9. doi: 10.1021/bi982246a.

Grosjean H, Gaspin C, Marck C, Decatur WA, de Crécy-Lagard V. RNomics and Modomics in the halophilic archaea Haloferax volcanii: identification of RNA modification genes. BMC Genomics. 2008 Oct 9;9:470. doi: 10.1186/1471-2164-9-470.

Hein N, Hannan KM, George AJ, Sanij E, Hannan RD. The nucleolus: an emerging target for cancer therapy. Trends Mol Med. 2013 Nov;19(11):643–54. doi: 10.1016/j.molmed.2013.07.005.

Helm M. Post-transcriptional nucleotide modification and alternative folding of RNA. Nucleic Acids Res. 2006 Feb 1;34(2):721–33. doi: 10.1093/nar/gkj471.

Hernández-Cid A, Lozano-Aponte J, Scior T. Molecular Dynamics and Docking Simulations of Homologous RsmE Methyltransferases Hints at a General Mechanism for Substrate Release upon Uridine Methylation on 16S rRNA. Int J Mol Sci. 2023 Nov 24;24(23):16722. doi: 10.3390/ijms242316722.

Hori H. Transfer RNA methyltransferases with a SpoU-TrmD (SPOUT) fold and their modified nucleosides in tRNA. Biomolecules. 2017 Feb 28;7(1):23. doi: 10.3390/biom7010023.

Jüttner M, Weiß M, Ostheimer N, Reglin C, Kern M, Knüppel R, Ferreira-Cerca S. A versatile cis-acting element reporter system to study the function, maturation and stability of ribosomal RNA mutants in archaea. Nucleic Acids Res. 2020 Feb 28;48(4):2073–2090. doi: 10.1093/nar/gkz1156.

Kirpekar F, Hansen LH, Rasmussen A, Poehlsgaard J, Vester B. The archaeon Haloarcula marismortui has few modifications in the central parts of its 23S ribosomal RNA. J Mol Biol. 2005 May 6;348(3):563–73. doi: 10.1016/j.jmb.2005.03.009.

Kiss-László Z, Henry Y, Bachellerie JP, Caizergues-Ferrer M, Kiss T. Site-specific ribose methylation of preribosomal RNA: a novel function for small nucleolar RNAs. Cell. 1996 Jun 28;85(7):1077–88. doi: 10.1016/s0092-8674(00)81308-2.

Lapeyre B, Purushothaman SK. Spb1p-directed formation of Gm2922 in the ribosome catalytic center occurs at a late processing stage. Mol Cell. 2004 Nov 19;16(4):663–9. doi: 10.1016/j.molcel.2004.10.022.

Lázaro E, Rodriguez-Fonseca C, Porse B, Ureña D, Garrett RA, Ballesta JP. A sparsomycin-resistant mutant of Halobacterium salinarium lacks a modification at nucleotide U2603 in the peptidyl transferase centre of 23 S rRNA. J Mol Biol. 1996 Aug 16;261(2):231–8. doi: 10.1006/jmbi.1996.0455.

Lee CH, Kiparaki M, Blanco J, Folgado V, Ji Z, Kumar A, Rimesso G, Baker NE. A Regulatory Response to Ribosomal Protein Mutations Controls Translation, Growth, and Cell Competition. Dev Cell. 2018 Aug 20;46(4):456–469.e4. doi: 10.1016/j.devcel.2018.07.003.

Liu B, Liang XH, Piekna-Przybylska D, Liu Q, Fournier MJ. Mis-targeted methylation in rRNA can severely impair ribosome synthesis and activity. RNA Biol. 2008 Oct-Dec;5(4):249–54. doi: 10.4161/rna.6916.

Liu J, Xu Y, Stoleru D, Salic A. Imaging protein synthesis in cells and tissues with an alkyne analog of puromycin. Proc Natl Acad Sci U S A. 2012 Jan 10;109(2):413–8. doi: 10.1073/pnas.1111561108.

Londei P, Ferreira-Cerca S. Ribosome Biogenesis in Archaea. Front Microbiol. 2021 Jul 22;12:686977. doi: 10.3389/fmicb.2021.686977.

Maden BE. The numerous modified nucleotides in eukaryotic ribosomal RNA. Prog Nucleic Acid Res Mol Biol. 1990;39:241–303. doi: 10.1016/s0079-6603(08)60629-7.

Mahto SK, Chow CS. Probing the stabilizing effects of modified nucleotides in the bacterial decoding region of 16S ribosomal RNA. Bioorg Med Chem. 2013 May 15;21(10):2720–6. doi: 10.1016/j.bmc.2013.03.010.

Marygold SJ, Roote J, Reuter G, Lambertsson A, Ashburner M, Millburn GH, Harrison PM, Yu Z, Kenmochi N, Kaufman TC, Leevers SJ, Cook KR. The ribosomal protein genes and Minute loci of Drosophila melanogaster. Genome Biol. 2007;8(10):R216. doi: 10.1186/gb-2007-8-10-r216.

Maxwell ES, Fournier MJ. The small nucleolar RNAs. Annu Rev Biochem. 1995;64:897–934. doi: 10.1146/annurev.bi.64.070195.004341.

Michel G, Sauvé V, Larocque R, Li Y, Matte A, Cygler M. The structure of the RlmB 23S rRNA methyltransferase reveals a new methyltransferase fold with a unique knot. Structure. 2002 Oct;10(10):1303–15. doi: 10.1016/s0969-2126(02)00852-3.

Misra VK, Draper DE. On the role of magnesium ions in RNA stability. Biopolymers. 1998;48(2-3):113–35. doi: 10.1002/(SICI)1097-0282(1998)48:2<113::AID-BIP3>3.0.CO;2-Y.

Nierhaus KH. Mg2+, K+, and the ribosome. J Bacteriol. 2014 Nov;196(22):3817–9. doi: 10.1128/JB.02297-14.

Noon KR, Bruenger E, McCloskey JA. Posttranscriptional modifications in 16S and 23S rRNAs of the archaeal hyperthermophile Sulfolobus solfataricus. J Bacteriol. 1998 Jun;180(11):2883–8. doi: 10.1128/JB.180.11.2883-2888.1998.

Petrov AS, Bernier CR, Hsiao C, Okafor CD, Tannenbaum E, Stern J, Gaucher E, Schneider D, Hud NV, Harvey SC, Williams LD. RNA-magnesium-protein interactions in large ribosomal subunit. J Phys Chem B. 2012 Jul 19;116(28):8113–20. doi: 10.1021/jp304723w.

Piekna-Przybylska D, Decatur WA, Fournier MJ. The 3D rRNA modification maps database: with interactive tools for ribosome analysis. Nucleic Acids Res. 2008 Jan;36(Database issue):D178–83. doi: 10.1093/nar/gkm855.

Pletnev P, Guseva E, Zanina A, Evfratov S, Dzama M, Treshin V, Pogorel’skaya A, Osterman I, Golovina A, Rubtsova M, Serebryakova M, Pobeguts OV, Govorun VM, Bogdanov AA, Dontsova OA, Sergiev PV. Comprehensive Functional Analysis of Escherichia coli Ribosomal RNA Methyltransferases. Front Genet. 2020 Feb 27;11:97. doi: 10.3389/fgene.2020.00097.

Polacek N, Mankin AS. The ribosomal peptidyl transferase center: structure, function, evolution, inhibition. Crit Rev Biochem Mol Biol. 2005 Sep-Oct;40(5):285–311. doi: 10.1080/10409230500326334.

Porse BT, Thi-Ngoc HP, Garrett RA. The donor substrate site within the peptidyl transferase loop of 23 S rRNA and its putative interactions with the CCA-end of N-blocked aminoacyl-tRNA(Phe). J Mol Biol. 1996 Dec 6;264(3):472–83. doi: 10.1006/jmbi.1996.0655.

Qi X, Wang W, Dong H, Liang Y, Dong C, Ly H. Expression and X-Ray Structural Determination of the Nucleoprotein of Lassa Fever Virus. Methods Mol Biol. 2018;1604:179–188. doi: 10.1007/978-1-4939-6981-4_12.

Saebøe-Larssen S, Lyamouri M, Merriam J, Oksvold MP, Lambertsson A. Ribosomal protein insufficiency and the minute syndrome in Drosophila: a dose-response relationship. Genetics. 1998 Mar;148(3):1215–24. doi: 10.1093/genetics/148.3.1215.

Schapira M. Structural Chemistry of Human RNA Methyltransferases. ACS Chem Biol. 2016 Mar 18;11(3):575–82. doi: 10.1021/acschembio.5b00781.

Sergiev PV, Aleksashin NA, Chugunova AA, Polikanov YS, Dontsova OA. Structural and evolutionary insights into ribosomal RNA methylation. Nat Chem Biol. 2018 Feb 14;14(3):226–235. doi: 10.1038/nchembio.2569.

Sharma S, Yang J, Düttmann S, Watzinger P, Kötter P, Entian KD. Identification of novel methyltransferases, Bmt5 and Bmt6, responsible for the m3U methylations of 25S rRNA in Saccharomyces cerevisiae. Nucleic Acids Res. 2014 Mar;42(5):3246–60. doi: 10.1093/nar/gkt1281. Epub 2013 Dec 11.

Shen F, Huang W, Huang JT, Xiong J, Yang Y, Wu K, Jia GF, Chen J, Feng YQ, Yuan BF, Liu SM. Decreased N(6)-methyladenosine in peripheral blood RNA from diabetic patients is associated with FTO expression rather than ALKBH5. J Clin Endocrinol Metab. 2015 Jan;100(1):E148–54. doi: 10.1210/jc.2014-1893.

Sirum-Connolly K, Mason TL. Functional requirement of a site-specific ribose methylation in ribosomal RNA. Science. 1993 Dec 17;262(5141):1886–9. doi: 10.1126/science.8266080.

Sloan KE, Warda AS, Sharma S, Entian KD, Lafontaine DLJ, Bohnsack MT. Tuning the ribosome: The influence of rRNA modification on eukaryotic ribosome biogenesis and function. RNA Biol. 2017 Sep 2;14(9):1138–1152. doi: 10.1080/15476286.2016.1259781.

Strassler SE, Bowles IE, Dey D, Jackman JE, Conn GL. Tied up in knots: Untangling substrate recognition by the SPOUT methyltransferases. J Biol Chem. 2022 Oct;298(10):102393. doi: 10.1016/j.jbc.2022.102393.

Streit D, Schleiff E. The Arabidopsis 2’-O-Ribose-Methylation and Pseudouridylation Landscape of rRNA in Comparison to Human and Yeast. Front Plant Sci. 2021 Jul 26;12:684626. doi: 10.3389/fpls.2021.684626.

Taoka M, Nobe Y, Yamaki Y, Sato K, Ishikawa H, Izumikawa K, Yamauchi Y, Hirota K, Nakayama H, Takahashi N, Isobe T. Landscape of the complete RNA chemical modifications in the human 80S ribosome. Nucleic Acids Res. 2018 Oct 12;46(18):9289–9298. doi: 10.1093/nar/gky811.

Taylor AB, Meyer B, Leal BZ, Kötter P, Schirf V, Demeler B, Hart PJ, Entian KD, Wöhnert J. The crystal structure of Nep1 reveals an extended SPOUT-class methyltransferase fold and a pre-organized SAM-binding site. Nucleic Acids Res. 2008 Mar;36(5):1542–54. doi: 10.1093/nar/gkm1172.

Tkaczuk KL, Dunin-Horkawicz S, Purta E, Bujnicki JM. Structural and evolutionary bioinformatics of the SPOUT superfamily of methyltransferases. BMC Bioinformatics. 2007 Mar 5;8:73. doi: 10.1186/1471-2105-8-73.

Treiber T, Treiber N, Plessmann U, Harlander S, Daiß JL, Eichner N, Lehmann G, Schall K, Urlaub H, Meister G. A Compendium of RNA-Binding Proteins that Regulate MicroRNA Biogenesis. Mol Cell. 2017 Apr 20;66(2):270–284.e13. doi: 10.1016/j.molcel.2017.03.014.

van Buul CP, Visser W, van Knippenberg PH. Increased translational fidelity caused by the antibiotic kasugamycin and ribosomal ambiguity in mutants harbouring the ksgA gene. FEBS Lett. 1984 Nov 5;177(1):119–24. doi: 10.1016/0014-5793(84)80994-1.

van Tran N, Ernst FGM, Hawley BR, Zorbas C, Ulryck N, Hackert P, Bohnsack KE, Bohnsack MT, Jaffrey SR, Graille M, Lafontaine DLJ. The human 18S rRNA m6A methyltransferase METTL5 is stabilized by TRMT112. Nucleic Acids Res. 2019 Sep 5;47(15):7719–7733. doi: 10.1093/nar/gkz619.

Wang T, Birsoy K, Hughes NW, Krupczak KM, Post Y, Wei JJ, Lander ES, Sabatini DM. Identification and characterization of essential genes in the human genome. Science. 2015 Nov 27;350(6264):1096–101. doi: 10.1126/science.aac7041.

Yang J, Sharma S, Watzinger P, Hartmann JD, Kötter P, Entian KD. Mapping of Complete Set of Ribose and Base Modifications of Yeast rRNA by RP-HPLC and Mung Bean Nuclease Assay. PLoS One. 2016 Dec 29;11(12):e0168873. doi: 10.1371/journal.pone.0168873.

Youngman EM, Brunelle JL, Kochaniak AB, Green R. The active site of the ribosome is composed of two layers of conserved nucleotides with distinct roles in peptide bond formation and peptide release. Cell. 2004 May 28;117(5):589–99. doi: 10.1016/s0092-8674(04)00411-8.

Zhang H, Wan H, Gao ZQ, Wei Y, Wang WJ, Liu GF, Shtykova EV, Xu JH, Dong YH. Insights into the catalytic mechanism of 16S rRNA methyltransferase RsmE (m³U1498) from crystal and solution structures. J Mol Biol. 2012 Nov 2;423(4):576–89. doi: 10.1016/j.jmb.2012.08.016.

